# split-intein Gal4 provides intersectional genetic labeling that is fully repressible by Gal80

**DOI:** 10.1101/2023.03.24.534001

**Authors:** Ben Ewen-Campen, Haojiang Luan, Jun Xu, Rohit Singh, Neha Joshi, Tanuj Thakkar, Bonnie Berger, Benjamin H. White, Norbert Perrimon

## Abstract

The split-Gal4 system allows for intersectional genetic labeling of highly specific cell-types and tissues in *Drosophila*. However, the existing split-Gal4 system, unlike the standard Gal4 system, cannot be repressed by Gal80, and therefore cannot be controlled temporally. This lack of temporal control precludes split-Gal4 experiments in which a genetic manipulation must be restricted to specific timepoints. Here, we describe a new split-Gal4 system based on a self-excising split-intein, which drives transgene expression as strongly as the current split-Gal4 system and Gal4 reagents, yet which is fully repressible by Gal80. We demonstrate the potent inducibility of “split-intein Gal4” *in vivo* using both fluorescent reporters and via reversible tumor induction in the gut. Further, we show that our split-intein Gal4 can be extended to the drug-inducible GeneSwitch system, providing an independent method for intersectional labeling with inducible control. We also show that the split-intein Gal4 system can be used to generate highly cell-type specific genetic drivers based on *in silico* predictions generated by single cell RNAseq (scRNAseq) datasets, and we describe a new algorithm (“Two Against Background” or TAB) to predict cluster-specific gene pairs across multiple tissue-specific scRNA datasets. We provide a plasmid toolkit to efficiently create split-intein Gal4 drivers based on either CRISPR knock-ins to target genes or using enhancer fragments. Altogether, the split-intein Gal4 system allows for the creation of highly specific intersectional genetic drivers that are inducible/repressible.

**Significance statement:** The split-Gal4 system allows *Drosophila* researchers to drive transgene expression with extraordinary cell type specificity. However, the existing split-Gal4 system cannot be controlled temporally, and therefore cannot be applied to many important areas of research. Here, we present a new split-Gal4 system based on a self-excising split-intein, which is fully controllable by Gal80, as well as a related drug-inducible split GeneSwitch system. This approach can both leverage and inform single-cell RNAseq datasets, and we introduce an algorithm to identify pairs of genes that precisely and narrowly mark a desired cell cluster. Our split-intein Gal4 system will be of value to the *Drosophila* research community, and allow for the creation of highly specific genetic drivers that are also inducible/repressible.

## Introduction

The ability to restrict transgene expression to specific, genetically-defined cell types using binary expression systems such as Gal4/UAS, LexA/LexAOP and QF/QUAS has profoundly transformed *Drosophila* research (1–3). In particular, the Gal4 system has been deployed extraordinarily effectively, with thousands of Gal4 drivers available in *Drosophila* resource centers. However, the lack of tissue- and cell-type specificity of many Gal4 drivers remains a drawback. This is especially true for certain areas of research. For example, studies of inter-organ communication in which a Gal4-driven manipulation is performed in one tissue and the effects are measured in a distant tissue must take special care to avoid the confounding effects of Gal4 expression outside of the intended tissue (4). Similarly, many neurobiological studies require Gal4 expression to be limited to one, or very few, transcriptionally-defined neurons, which is not generally possible using standard Gal4 drivers, even when driven by 2-3kb genomic enhancer fragments (5–7).

The split-Gal4 system was developed to overcome the issue of limited cell-type specificity, by restricting transgene expression to those cells that co-express two independent enhancers, a strategy termed “intersectional genetic labeling” (8, 9). In split-Gal4, the N-terminal 147 amino acids of Gal4, which includes its DNA binding domain (Gal4DBD) (10) and its dimerization domain (11), is expressed under the control of one enhancer, while a potent transcriptional activator domain (AD) from either VP16 or p65 is expressed under the control of a second enhancer (**Figure 1**) (8, 12). The Gal4DBD and the VP16/p65 activation domains are each flanked by a leucine zipper domain, which heterodimerize in any cell expressing both components, and reconstitute a functional Gal4-like transcription factor (8, 9). The split-Gal4 system has been successfully used to build thousands of exquisitely specific genetic drivers, especially in the *Drosophila* nervous system where split-Gal4 lines are now routinely utilized to drive expression in a single pair of neurons (6, 7), and in the adult gut, where thousands of split-Gal4 lines have been characterized (13, 14). The ability to create split-Gal4 lines that are specifically expressed in the same patterns as genes of interest using “trojan exons” or other knock-in strategies has further augmented the power of the split-Gal4 method (15). This capability has particular promise in permitting the construction of genetic driver lines that target transcriptionally distinct clusters identified via scRNAseq studies (16). For such clusters, the intersection of at least two genetic markers is typically necessary to uniquely identify specific clusters.

**Figure 1.**
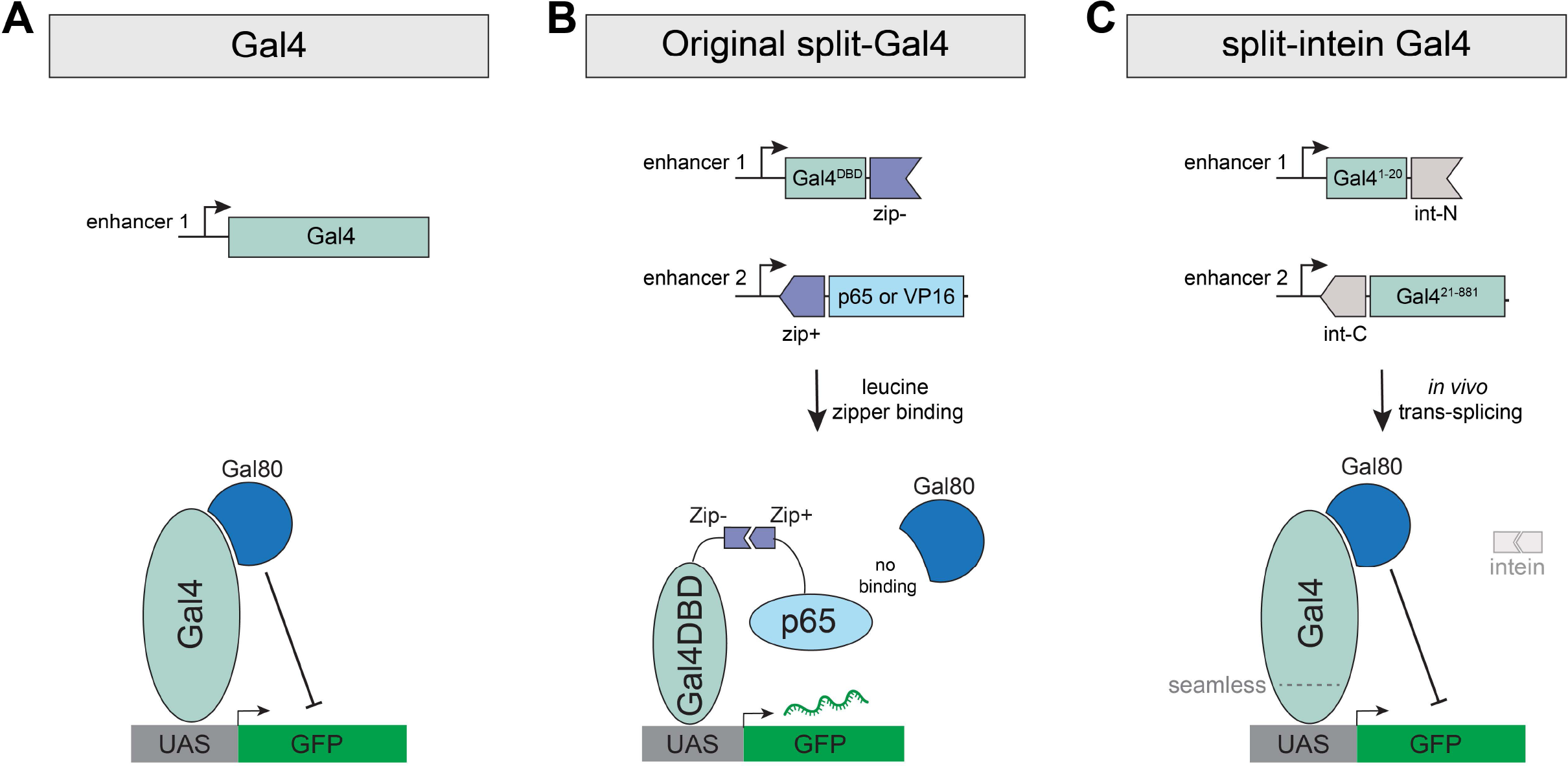
Schematic comparison of Gal4, Split-Gal4, and split-intein Gal4. (A) The Gal4 transcription factor binds to UAS sequences to drive transcription, and can be repressed by the binding of Gal80. Gal4 is drawn here as a monomer, but functions as a dimer *in vivo.* (B) In the original split-Gal4 system, the Gal4DBD and a strong transcriptional activator (VP16 or p65) are each driven by separate enhancers, and reconstituted in cells by leucine zipper domains. Gal80 cannot bind or repress the split-Gal4 complex. (C) In the split-intein Gal4 system, two fragments of the Gal4 protein, each flanked by a split-intein, are independently driven by separate enhancers, and seamlessly trans-spliced to reconstitute a functional, wildtype Gal4 protein, which can be repressed by Gal80.

But while the split-Gal4 system has effectively solved the problem of restricting expression in anatomical space, the existing split-Gal4 system cannot be controlled in time using any available technique. This is in contrast to standard Gal4 drivers, which can be temporally controlled using a temperature sensitive variant of the Gal80 repressor (Gal80^ts^) (17). By shifting between a permissive temperature (18°C), where Gal80^ts^ fully represses Gal4 expression, and a restrictive temperature (29°C), where Gal80^ts^ is inactivated and Gal4 becomes active, researchers can restrict genetic manipulations to specific time periods or developmental stages. By contrast to the standard Gal4 system, the split-Gal4 system is completely insensitive to the Gal80 repressor (9). This is because the region of Gal4 that is bound by Gal80, the C-terminal 30 amino acids (18), falls squarely within the Gal4 AD domain, which is replaced in existing split-Gal4 implementations with either VP16 or p65 in order to drive sufficiently high levels of expression (**Figure 1** and **2A**.) Thus, the Gal4DBD-VP16 or Gal4DBD-p65 protein complexes do not contain any binding site for Gal80 and therefore cannot be repressed (**Figure 1**).

**Figure 2.**
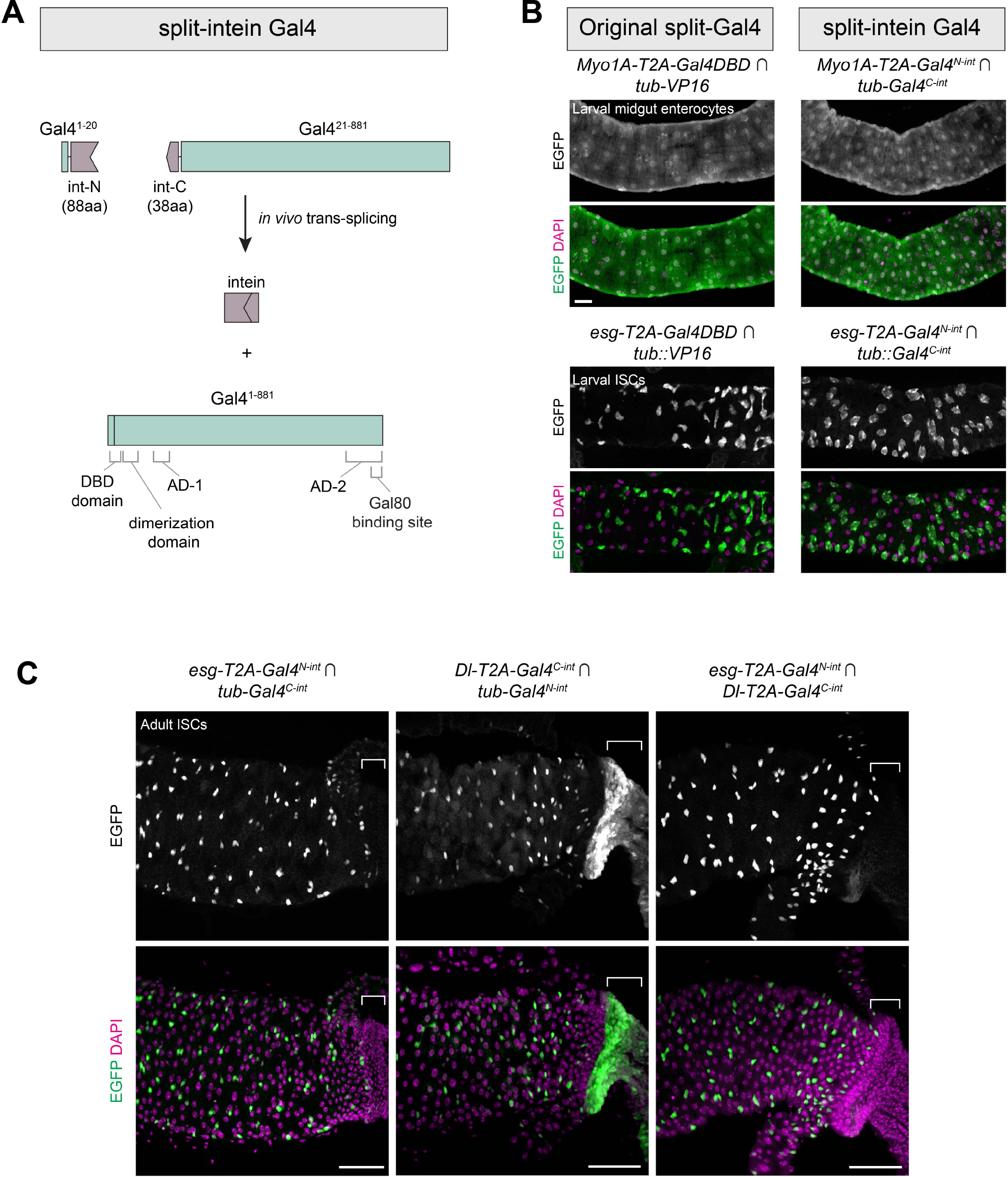
Split-intein Gal4 system drives intersectional expression at levels indistinguishable from split-Gal4. (A) Schematic diagram of the split-intein Gal4 system. Gal4 and gp-41 are drawn to scale, illustrating the N terminal DNA binding domain (DBD) and dimerization domain, and the overlap of the second activation domain (AD-2) with the Gal80 binding site. (B) Components for original split-Gal4 (left) or split-intein Gal4 (right) were knocked into two gut cell-type markers: *Myo1A,* which labels ECs, and *esg,* which labels ISCs. These knock-in lines were crossed to ubiquitously-expressed tester lines to visualize their full expression pattern. (C) Intersectional labeling of midgut intestinal stem cells using *esg* ∩ *Dl* split-intein Gal4 knock-in lines. Brackets indicate expression in the anterior hindgut which is driven by *Dl* but not *esg*, which is absent in their intersection. Anterior is to the left. scale bars = 50 µm.

There is thus a clear need for an intersectional labeling system that is repressible by Gal80^ts^ or otherwise inducible. Here, we describe two such systems. The first we term “split-intein Gal4.” This system combines the enhanced spatial control offered by split-Gal4 with the ability to strictly limit genetic manipulations to specific periods of time using existing Gal80^ts^ reagents. We demonstrate that split-intein Gal4 drives UAS transgenes at expression levels that are indistinguishable from existing split-Gal4 and Gal4 reagents and that it can be fully repressed by standard Gal80 reagents. The second system is a closely related drug-inducible GeneSwitch technique (“split-intein GeneSwitch”), which provides an alternative means to induce intersectional genetic labeling using a drug rather than a temperature shift. Finally, we demonstrate that the split-intein Gal4 system can be effectively used with scRNAseq datasets to generate split-intein Gal4 driver lines. To facilitate production of such lines, we present an algorithm to select gene pairs with low levels of predicted co-expression outside the cluster-of-interest. We provide a simple cloning and transgenesis workflow that can be used to generate large numbers of split intein-Gal4 lines, either via CRISPR-based knock-in or using enhancer fragments.

## Results

### Designing a split-Gal4 technique that can be repressed by Gal80

We wished to create an inducible/repressible intersectional labeling technique that can be controlled temporally. We focused our efforts on modifying the highly effective split-Gal4 concept, to make a split system that did not rely on leucine zipper heterodimerization and that incorporated the native Gal4 activation domain, thus rendering it sensitive to the Gal80^ts^ repressor. We devised two independent strategies, which we refer to as “split-intein Gal4” and “NanoTag split-Gal4.”

*Split-intein Gal4.* Native split-inteins consist of N- and C-terminal peptides that are fused to proteins encoded at separate genomic loci. Upon translation, these peptides associate with one another, self-excise, and seamlessly trans-splice the two adjacent polypeptide chains to which they are fused (19). Split inteins have been successfully exploited to generate split proteins used in other expression systems (20, 21), and we sought to use them here to reconstitute wildtype Gal4 from two functionally inert fragments.

In a series of pilot experiments in S2 cells, we tested three different cysteine residues to split Gal4 into two non-functional fragments, and four different split-intein systems (see *Methods* for full description) and ultimately identified the most potent system, which we refer to as “split-intein Gal4.” In this system, the Gal4 protein is split into two fragments: an N-terminal 20 amino acid portion (Gal4^N-int^) and the remaining C-terminal 861 amino acids (Gal4^C-int^), each flanked by components of the highly active *gp41-1* split-intein sequence (19) (**Figure 1**, **Figure 2A**). When these two fragments are co-expressed in a cell, the split intein activity is predicted to reconstitute the full wildtype Gal4 protein, which should be repressible by Gal80 (**Figure 1**, **Figure 2A**.) Previous studies in *C. elegans* have demonstrated a related approach, in which a DNA binding domain and an activator domain are trans-spliced via *gp41-1* split-intein (20). However, in that approach, a VP64 activator domain is used instead of the native Gal4 domains, and thus this system is not repressible by Gal80.

*NanoTag split-Gal4*. “NanoTags” are short epitope tags (<25 amino acids) that are recognized with very high affinity by single domain nanobodies. Recently, two high-affinity NanoTags, 127D01 and VHH05, have been adapted for a variety of applications *in vivo* in *Drosophila* (22). We designed a split-Gal4 system based on the affinity of the 127D01 tag and its genetically-encoded nanobody, Nb127D01. In a series of pilot experiments in S2R+ cells, we observed that Gal4DBD-Nb127D01 combined with Gal4AD-1×127D01 drove only very weak expression. However, when we fused three Nb12701 nanobodies in tandem to a Gal4DBD domain (Gal4DBD-3×Nb127D01), making it capable of recruiting three Gal4-AD molecules to each Gal4DBD domain, we observed robust transgene expression. We refer to this combination as NanoTag split-Gal4.

### Both split-intein Gal4 and NanoTag split-Gal4 function in *Drosophila* cell culture

To test the transcriptional activation strength of each system in cell culture, we transiently transfected either split-intein components (Gal4^N-int^ and Gal4^C-int^) or NanoTag Split-Gal4 components (Gal4DBD-3×Nb12701 and Gal4AD-1×127D01), all driven by a constitutive *actin* promoter, into S2R+ cells, along with a UAS:GFP reporter. As positive controls, we transfected full length Gal4, and standard split-Gal4 components, Zip-Gal4DBD and p65-Zip. Two days after transient transfection, we observed strong GFP expression for both split-intein Gal4 and NanoTag split-Gal4, at similar levels to Gal4 itself or to the existing split-Gal4 system (**Figure S1**).

We tested whether split-intein Gal4 and NanoTag Gal4 were repressible by Gal80 in S2R+ cells by co-transfecting these components with pTub:Gal80. As expected, wildtype Gal4, but not the existing split-Gal4 system (Gal4DBD-p65), was strongly repressed by Gal80 (**Figure S1**). Strikingly, both split-intein Gal4 and NanoTag split-Gal4 exhibited strong repression by Gal80, albeit slightly weaker than that observed for wildtype Gal4 (**Figure S1**). We conclude that both approaches offer robust transcriptional activation at levels similar to the existing split-Gal4 system, but have the critical advantage that they are also sensitive to the Gal80 repressor.

While both approaches showed promise in S2R+ cells, the 3:1 stoichiometry of Gal4AD:Gal4DBD in the NanoTag system suggested that this approach might require higher levels of Gal80 than the split-intein Gal4 approach to achieve the same level of repression. Since Gal80 expression levels *in vivo* will generally vary from cell-type to cell-type for any given Gal80 line and high sensitivity is therefore desirable, we chose to focus on the split-intein Gal4 system for additional *in vivo* testing.

### The split-intein Gal4 system activates high levels of intersectional UAS-driven expression *in vivo*

In order to be a broadly useful tool *in vivo*, split-intein Gal4 must meet three criteria. It must: (1) *drive robust expression in vivo*, at levels similar to existing split-Gal4 or Gal4 lines; (2) *drive clean intersectional labeling* that is not “leaky,” and includes only those cells expressing both Gal4^N-int^ and Gal4^C-int^ components; (3) *be fully repressible* using existing Gal80^ts^ lines.

To characterize split-intein Gal4 *in vivo*, we used CRISPR-mediated knock-in transgenesis to insert split-intein components into various genes with well-characterized expression patterns. We first selected two genes expressed in specific cell types of the midgut: *Myo1A* (aka *Myo31DF*) which is expressed in enterocytes (ECs)(23), and *esg,* which is expressed in intestinal stem cells (ISCs)(24). To permit direct comparison with the current split-Gal4 system, we also generated knock-ins of ZipGal4DBD into the same positions within *esg* and *Myo1A*. To create these knock-ins, we adapted the “drop-in” cloning method (25) to generate homology-driven repair (HDR) donor plasmids that would insert an in-frame T2A sequence, followed by the split-intein Gal4 or split-Gal4 component, into an early exon of the target gene. We also generated ubiquitously-expressed split-intein Gal4 components, driven by the *alphaTubulin48B* promoter, to use as “tester” lines to visualize the complete expression pattern of each knock-in.

We crossed *Myo1A-T2A-Gal4^N-int^* to the *tub-Gal4^C-int^; UAS:2×EGFP* tester line, and observed cell-type specific expression in larval ECs, at levels statistically indistinguishable from the original split-Gal4 system (mean pixel intensity measured in *n =* 4 larval guts; t(6)=0.9325, *p* = 0.3871) (**Figure 2B**). Similarly, *esg-T2A-Gal4^N-int^* ∩ *tub-Gal4^C-int^* (hereafter we follow the convention of using the ∩ symbol to indicate intersectional labeling) drove specific expression in ISCs at similar levels to the standard split-Gal4 system (*n =* 3 larval guts; t(4) = 2.22; *p* = 0.091) (**Figure 2B**). These results indicate that the split-intein Gal4 system functions robustly *in vivo*. Expression in the midgut was specific for the two targeted cell types, indicating that the Gal4^C-int^ fragment did not support leaky expression.

We then tested whether the split-intein Gal4 approach would successfully drive intersectional expression using two cell-type-specific knock-in lines. As *esg* and *Myo1A* are not co-expressed in the gut, we knocked *Gal4^C-int^* into the *Delta* (*Dl*) gene, which is also expressed in ISCs (26). As expected, *esg* ∩ *Dl* expression was observed in adult ISCs (**Figure 2B**). Importantly, the expression of *Dl* in the anterior hindgut was not observed in the *esg* ∩ *Dl* intersection, providing additional evidence that the split-intein system is not leaky (**Figure 2B**).

Thus, the split-intein Gal4 system satisfies the first two criteria identified above: it drives expression at similar levels to the existing split-Gal4 system, and expression can only be detected in cells co-expressing both components.

### Split-intein Gal4 is fully repressible by Gal80^ts^

The split-intein Gal4 system should seamlessly reconstitute wildtype Gal4, which, unlike the original split-Gal4 system, is repressible by Gal80 (**Figure 3A**). To confirm this is the case, we generated larvae expressing both *tub-*Gal4^N-int^ and *tub-*Gal4^C-int^ as well as *tub-Gal80^ts^*and a UAS:2×-EGFP reporter.

**Figure 3.**
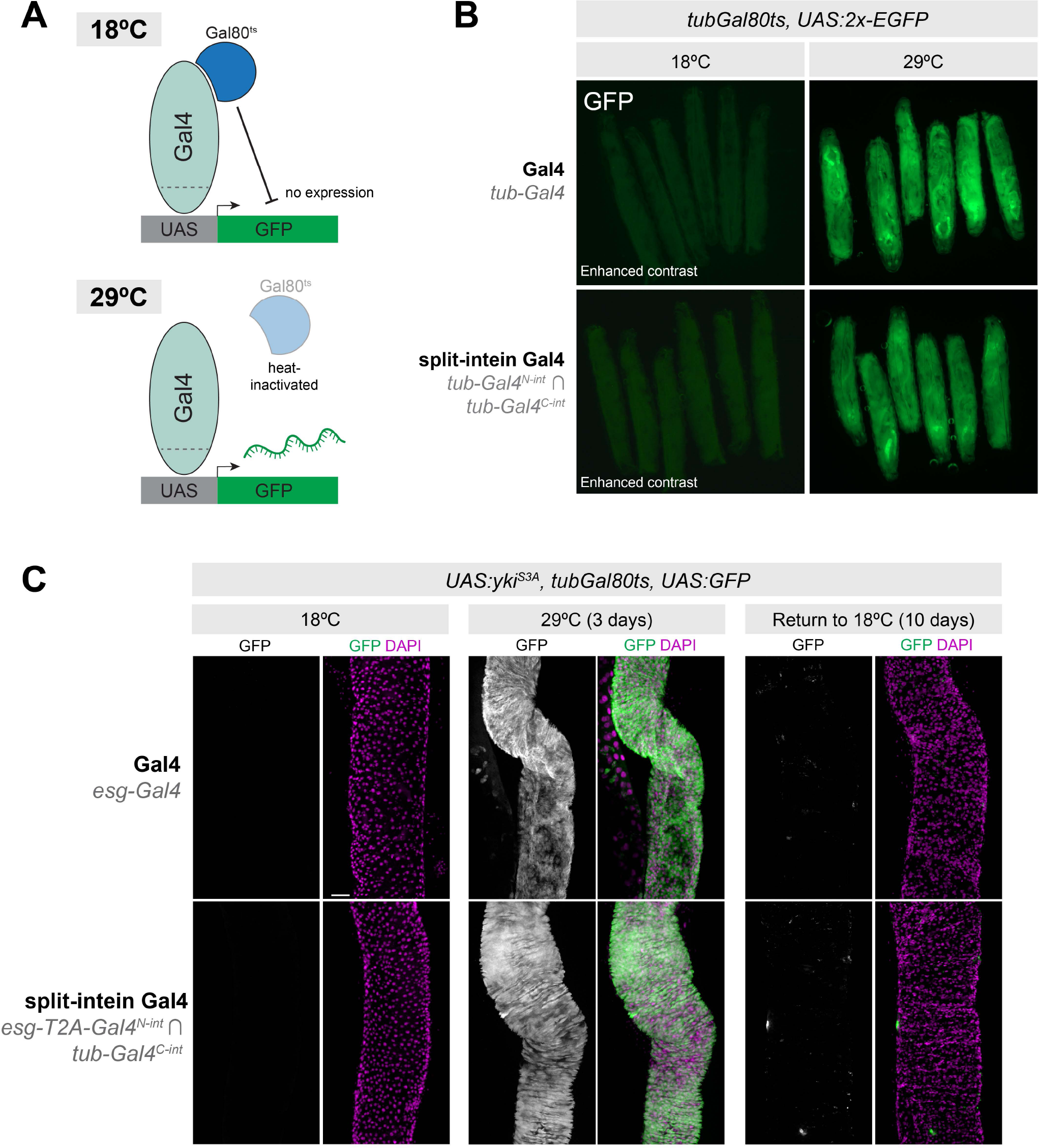
Split-intein Gal4 is fully repressible by Gal80. (A) Cartoon diagram of temperature inducible expression using Gal80^ts^. (B) *tub-Gal4^N-int^* ∩ *tub-Gal4^C-int^ tub-Gal80^ts^* > *UAS:2×EGFP* expression is fully repressed at 18°C and highly active at 29°C, as is *tub-*Gal4, *tub-*Gal80^ts^ > UAS:2×EGFP. Note that the 18°C images are shown with increased gain relative to the 29°C images in order to visualize the presence of larva. (C) Recapitulation of a well-characterized adult stem cell tumor system using split-intein Gal4. *esg-Gal4^N-int^ ∩ tub-Gal4^C-int^, tub-*Gal80^ts^ > UAS:*yki^S3A^* in adult ISCs, can be fully repressed throughout development and adult stages by *tubGal80^ts^*, and reversibly activated using temperature shift to 29°C. Anterior is up, scale bar in (C) is 50µm.

When *tub-Gal4^N-int^* ∩ *tub-Gal4^C-int^, tub-Gal80ts* > *UAS:2×EGFP* larva were grown at 18°C, no EGFP could be detected, similar to standard *tub-Gal4, tub-Gal80^ts^ > UAS:2×EGFP* larvae (**Figure 3B, left**) However, when grown at 29°C, strong EGFP expression was observed (**Figure 2B, right**). We also confirmed that, in the absence of Gal80^ts^, the split-intein Gal4 system does indeed drive EGFP expression at 18°C (**Figure S2**), indicating that the lack of EGFP expression at 18°C is not the result of compromised split intein trans-splicing, and is indeed due to Gal80 repression. These results also demonstrated that, like wildtype Gal4, split-intein Gal4 activity increases with temperature (27) (**Figure S2**). Together, these results demonstrate that split-intein Gal4 is repressible by Gal80.

To confirm that the Gal80 repression of split-intein Gal4 is sufficiently potent to fully repress strong, dominant phenotypes at 18°C, we turned to a widely used tumor model in the adult gut. When activated *yki* is expressed in ISCs using *esg*-Gal4, it generates severe tumor phenotypes in the adult gut (28–30). We used the split-intein Gal4 system to drive activated *yki* (*UAS:yki^S3A^*) in adult ISCs (*esg*-Gal4^N-int^ ∩ *tub-*Gal4^C-int^), in the presence of *tub-Gal80^ts^.* Grown at 18°C, these flies developed and eclosed normally, and displayed no EGFP expression or tumor growth in the ISCs, similar to the corresponding Gal4 system (**Figure 3C, left**). However, after a 3-day temperature shift, we observed a dramatic tumor phenotype indistinguishable from those produced by *esg-Gal4* (**Figure 3C, middle**). Further, we could re-repress this phenotype and the associated EGFP expression by shifting these flies back to 18°C for 10 days (**Figure 3C, right**). Altogether, these experiments confirm that split-intein Gal4 drives high levels of expression, is not leaky, and is fully repressible by existing Gal80^ts^ reagents.

The gene-specific split-intein Gal4 drivers described above were generated using CRISPR-based knock-ins, in order to fully recapitulate the endogenous expression pattern of the target gene. However, many researchers may wish to create Gal80-sensitive split-intein Gal4 drivers using specific enhancer fragments, as has been done successfully for thousands of split-Gal4 lines that have been generated for the VT split-Gal4 collection (6, 7). To facilitate this approach, we modified the pBPZpGAL4DBD and pBPp65ADZp destination vectors (12) to encode split-intein Gal4 components. These vectors are compatible with the well-established Gateway LR-based cloning workflow for generating enhancer fragment-driven split-Gal4 vectors (12, 31). To demonstrate the effectiveness of this approach, we selected a genomic fragment known to drive expression in the adult gut ISCs, VT024642 (13), and cloned this fragment into our pBP-Gal4^N-int^ destination vector. As predicted, *VT024642-Gal4^N-int^* ∩ *tub-Gal4^C-int^* drove strong, specific expression in adult ISCs (**Figure S3**.)

### Split-intein Gal4 components can be adapted to GeneSwitch for drug-inducible intersectional labeling

Having established that split-intein Gal4 is highly effective, we reasoned that this system should also be adaptable to the drug-inducible GeneSwitch system (32). In GeneSwitch, the Gal4DBD (the first 93 amino acids of the Gal4 protein) is fused to an RU486-sensitive ligand binding domain (PR-LBD) and a p65 transcriptional activator domain. In the absence of RU486 (RU), the GeneSwitch complex is inactive, whereas in the presence of RU, the complex undergoes a conformational change that allows for the transcriptional activation of UAS-driven transgenes.

We noted that the Gal4^N-int^ fragment, compromising the first 20 amino acids of Gal4, could be compatible with the corresponding C-terminal region of GeneSwitch (Gal4^21-93^-PR-LBD-p65 aka GeneSwitch^C-int^) flanked by a split-intein (**Figure 4A**.) In other words, the same Gal4^N-int^ lines could be crossed to either a Gal4^C-int^ for split-intein Gal4 expression, or to a GeneSwitch^C-int^ line for split-intein GeneSwitch expression. To test this, we generated a transgenic line expressing split-intein-GeneSwitch^C-int^ under the control of the *tub* promoter. We crossed *tub-split-intein-GeneSwitch^C-int^; UAS:2×EGFP* to *esg-T2A-Gal4^N-int^,* and split the F1 adult flies into RU- and RU+ minus treatments for six days. In the absence of RU, we observed no GFP expression in the adult gut, whereas flies fed RU-containing food displayed strong and specific GFP in adult ISCs (**Figure 4B, top**). In parallel, we crossed *tub-split-intein-GeneSwitch^C-int^; AS:2×EGFP* to *esg-T2A-Gal4^N-int^; UAS:yki^S3A^* to test whether we could successfully regulate tumor growth via RU feeding. While RU- flies displayed no EGFP expression and wildtype gut morphology, RU+ flies displayed strong ISC tumor phenotypes resembling those observed using either Gal4 or split-intein Gal4 (**Figure 4B, bottom**). Thus, the split-intein GeneSwitch system successfully combines intersectional genetic labeling with the RU-inducibility of GeneSwitch.

**Figure 4.**
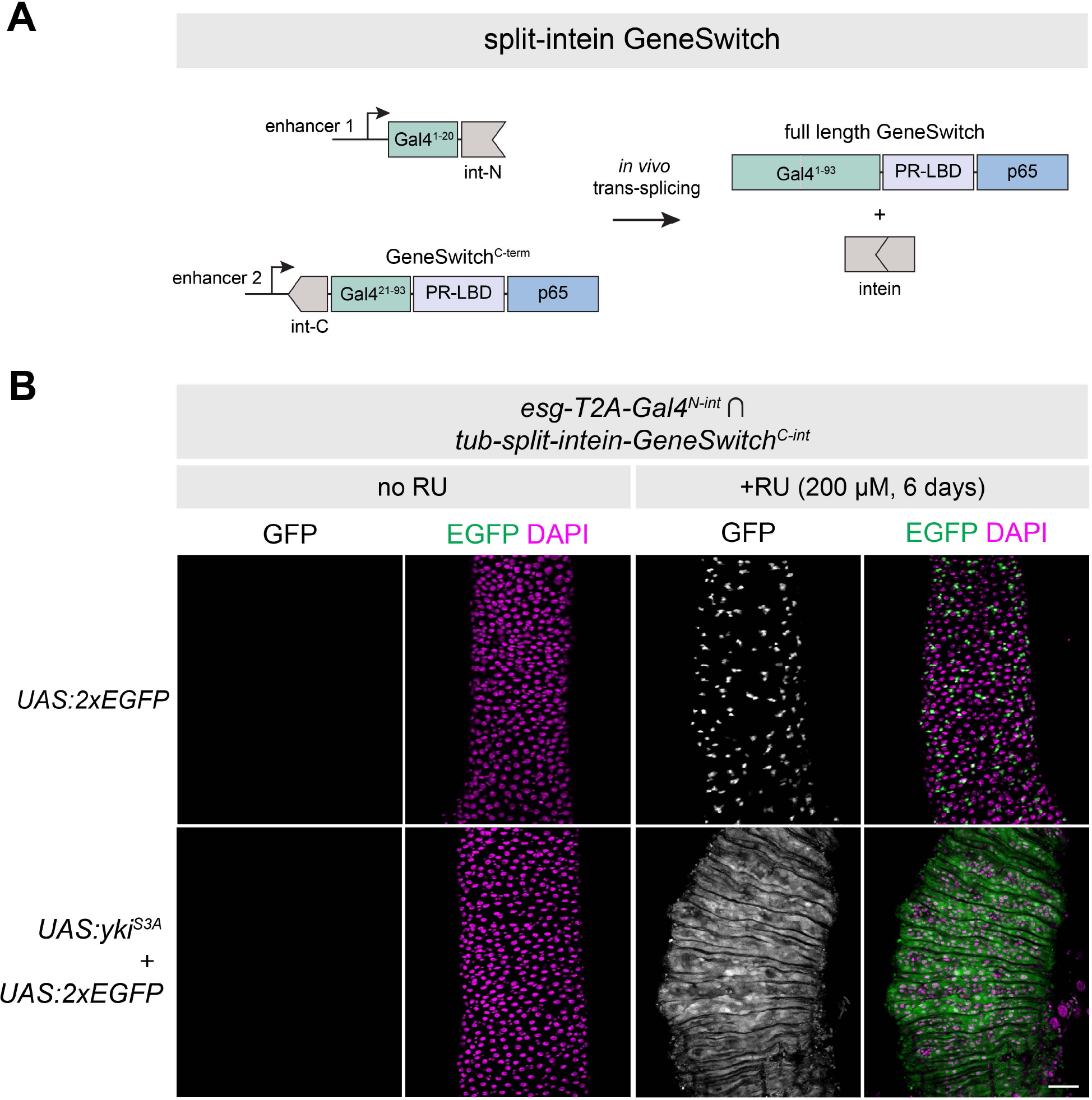
Drug-inducible intersectional labeling using split-intein GeneSwitch. (A) Cartoon schematic of the split-intein GeneSwitch system, not drawn to scale. The same N-terminal fragment of Gal4 used in split-intein Gal4 can be combined with the C-terminus of the GeneSwitch system, which includes amino acids 21-93 of Gal4, a progesterone ligand binding domain (PR-LBD) and the p65 transcriptional activator. (B) Drug-inducible ISC tumor model using the *esg*-Gal4^N-int^ *∩ tub-*GeneSwitch^C-int^ > UAS:*yki^S3A^*. Anterior is up. Scale bar = 50 µm.

Importantly, the GeneSwitch system has been shown to be leaky in some tissues, with detectable expression in the absence of RU (32–34). Given that split-intein GeneSwitch simply reconstitutes the existing GeneSwitch protein, we predicted this leaky expression would also be the case with split-intein GeneSwitch. We tested this by crossing the ubiquitously expressed *tub-split-intein-GeneSwitch^C-int^*tester line to three additional Gal4^N-int^ lines: *Myo1A-T2A-Gal4^N-int^*, *Dl-T2A-Gal4^N-int^*, and *tub-Gal4^N-int^*. While we observed clean RU-dependent expression in adult ISCs with both *esg* and *Dl* (**Figure S4A,B**), we observed leaky expression in a portion of the adult gut using *Myo1A* (**Figure S4C**), as well in the larval gut when using the *tub* promoter (**Figure S4D.**) Thus, as with existing GeneSwitch reagents, it will be important for researchers to carefully characterize the RU- and RU+ expression patterns for split-intein GeneSwitch lines.

### Mapping scRNA clusters to anatomy using split-intein Gal4 drivers

One particularly promising use of intersectional labeling techniques such as split-intein Gal4 is to characterize the many transcriptionally-defined “clusters” of cells that are identified using scRNAseq. Single cell and single nuclei transcriptomic atlases are now available for many individual *Drosophila* tissues, as well as for the entire adult body (35). These atlases identify many different distinct cell types within a given tissue based on transcriptional similarity, many of which remain uncharacterized either anatomically or functionally. In most cases, a single genetic marker is insufficiently specific to label a cluster, and a minimum of two co-expressed genes is generally required to demarcate a cluster (36). Thus, intersectional genetic labeling approaches are a promising tool to interrogate hypotheses generated via scRNAseq. The promise of this approach has recently been piloted in the *Drosophila* optic lobe (16).

To explore how the split-intein Gal4 system can leverage and inform scRNAseq datasets, we began with a recent atlas of the adult midgut, which identified 22 transcriptionally-distinct cell types (37). To pick pairs of genes that uniquely mark scRNA clusters, we implemented a recently described gene selection algorithm, NS-Forest version 2.0 (36). NS-Forest v2 is a machine learning algorithm that estimates the minimum number of marker genes that can be used to uniquely define scRNAseq clusters. Using NS-Forest v2 to guide gene pair selection, we generated transgenic split-intein Gal4 lines to mark three of these clusters *in vivo*: (1) aEC-3, predicted to be a subset of anterior enterocytes, marked by *Peritrophin-15a* ∩ *CG4830* (2) iron and copper cells, a functional analog of the human stomach located midway between the anterior and posterior of the midgut, marked by *CG43774* ∩ *thetaTry*; and (3) pEC-1, predicted to be a subset of posterior enterocytes, marked by *LManV* ∩ *ninaD* (**Figure 5**.)

**Figure 5.**
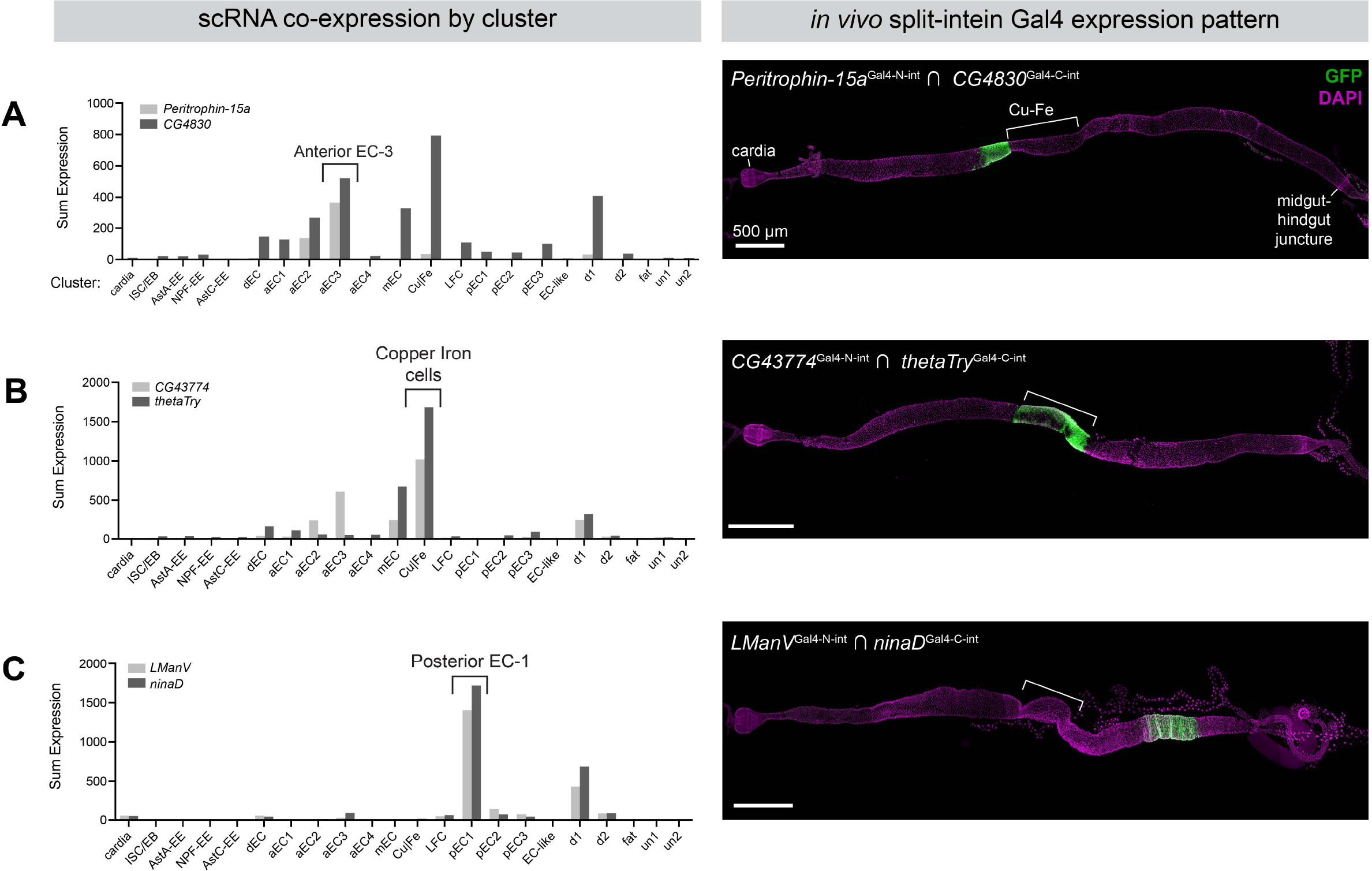
Mapping scRNA clusters to anatomy using split-intein Gal4 lines based on NS-Forest v2 predictions. Based on a gut-specific scRNA dataset from Hung *et al.* (2020), the NS-Forest v2 algorithm identified pairs of genes to mark specific clusters in the adult gut. Left: Summed expression in each of the 22 clusters is shown on the left. Right: *in vivo* expression of split-intein Gal4 lines in adult guts. Brackets indicate the approximate position of the copper and iron cell (Cu-Fe) region. (A) *Peritrophin-15a* ∩ *CG4830* drives expression in a band of enterocytes anterior to the copper cells. (B) *CG43774* ∩ *thetaTry* drives expression in the copper and iron cells. (C) *LManV* ∩ *ninaD* drives expression in a posterior band of enterocytes. Anterior is to the left.

We generated split-intein Gal4 lines for each of these three pairs, and examined their expression using UAS:2×EGFP. In each case, the expression pattern conformed well with the predicted location of the cell cluster. Our aEC3 split-intein Gal4 line *Peritrophin-15a* ∩ *CG4830* drove expression in a band of enterocytes anterior to the copper cells, corresponding to the “A3” region identified by (38) (**Figure 5A**). Similarly, the split-intein Gal4 line *CG43774* ∩ *thetaTry,* predicted to express in iron and copper cells, drove expression in precisely this region, and the pEC-1 line (**Figure 5B**). *LManV* ∩ *ninaD* drove expression in a band of enterocytes posterior to the copper cells (**Figure 5C**). Thus, using *in silico* predictions to guide gene selection, we were able to successfully use split-intein Gal4 to label anatomically-distinct populations of cells along the anterior-posterior axis of the adult midgut, demonstrating the potential power to identify and functionally characterize cell types identified via scRNAseq studies.

### Developing an algorithm to pick cluster-specific gene pairs with minimal co-expression across the “whole body” scRNA dataset

Subsequent to the publication of the midgut-specific scRNAseq dataset described by (37) and utilized above, the Fly Cell Atlas Consortium published scRNAseq datasets covering many additional adult *Drosophila* tissues, as well as a whole body dataset (35). As the set of potential off-target clusters now spans the whole body, it becomes more difficult to obtain gene pairs that mark a specific cluster. In particular, for each of the midgut-relevant gene pairs predicted by NS-Forest v2, we examined the *in silico* co-expression of these gene pairs across the entire body Fly Cell Atlas dataset. For the large majority of gene pairs, we observed that the NS-Forest v2 gene pairs had a high degree of co-expression in multiple other tissues. This has practical implications: it is crucial to identify intersectional genetic drivers that are exclusively expressed in specific tissues, with no additional expression anywhere else in the body.

We therefore sought to develop an algorithm that would specifically identify cluster-specific gene pairs that maximize co-expression in the cluster-of-interest, while minimizing additional co-expression in both the tissue-of-interest, as well as across other scRNAseq datasets from the same organism. To do so, we developed a gene-selection algorithm that we call “Two Against Background” or TAB.

The TAB algorithm is schematized in **Figure 6A**. Briefly, TAB incorporates three features to guide gene pair selection: first, it incorporates bulk RNA-seq data to supplement scRNA-seq expression estimates when available. Second, it emphasizes selecting genes with robust within-cluster expression profiles that are stable and not highly variable. Third, to calibrate the importance of cluster specificity relative to these robustness considerations, it employs a hyperparameter optimization approach by incorporating rankings by a researcher who is blinded to the parameters.

**Figure 6.**
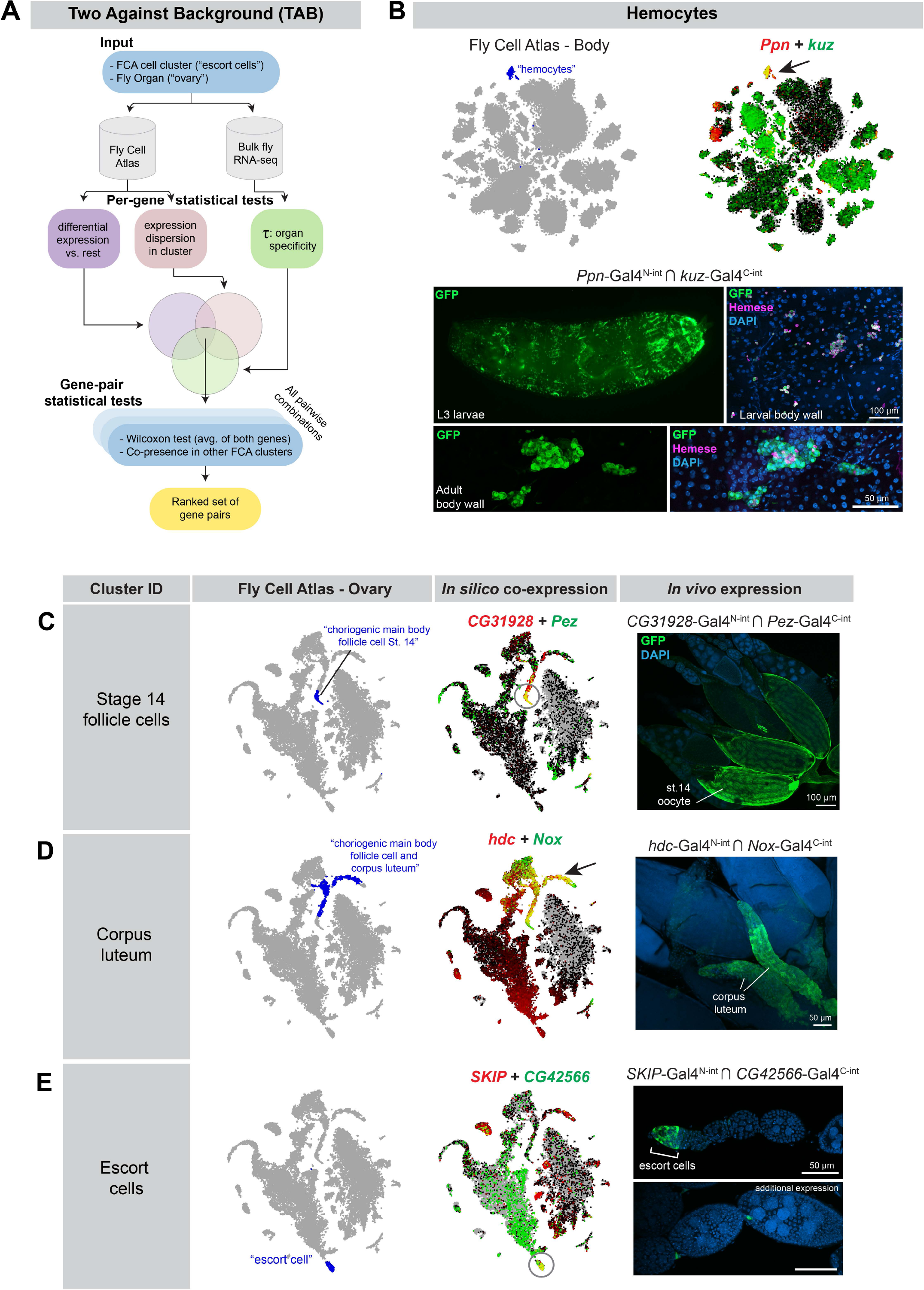
Characterization of split-intein Gal4 lines based on Two Against Background (TAB) algorithm predictions. (A) Schematic of the TAB algorithm to pick gene pairs that specifically mark scRNAseq clusters. (B) Hemocyte-specific split-intein Gal4 line based on TAB gene pairs. Top left: Fly Cell Atlas 10× “Body” atlas, with hemocyte cluster indicated in blue. Top right: *in silico* prediction of *Ppn* and *kuz* co-expression in the hemocyte cluster, with co-expression shown in yellow. Bottom: *In vivo* expression of the *Ppn* ∩ *kuz* split intein-Gal4 driver in larval and adult hemocytes. Hemocytes are co-stained with the H2 antibody against Hemese, a pan-hemocyte marker. (C-E) *In silico* predictions of co-expression in three different clusters from the FCA “Ovary” dataset, and *in vivo* co-expression in the indicated cell types of the ovary. scRNAseq data are screenshots from the FCA data viewer for the “10×, stringent” datasets.

As input, TAB requires both the cell cluster (e.g., “escort cells”) as well as the containing tissue (e.g. “ovary.”) To ensure that the genes being selected are well-expressed in the tissue, we cross-reference the gene’s bulk scRNA-seq expression levels in the corresponding organ. While the organ-level resolution is coarser than scRNA-seq cell clusters, the higher quality of bulk expression data is a valuable corrective for noisy scRNA-seq expression estimates. To quantify the specificity of any gene to the organ of interest, we use the Tau statistic. In addition, we require that the gene’s within-cluster expression be stable and not highly variable, i.e. the dispersion (variance/mean) of the gene expression in the cluster is limited.

In the TAB algorithm, a candidate set of gene pairs is created for each cluster-of-interest using the intersection of three metrics: the Tau statistic, dispersion metrics, and t-test of differentiation against all other clusters (**Figure 6A**). From this candidate set, we evaluate all pairwise combinations of genes and select pairs that are effective at distinguishing the cluster of interest from others. One of the metrics we consider is the number of other clusters where both the candidate genes are potential markers. We also introduce an additional metric: we construct a metagene as the average of the two genes, and perform the Wilcoxon rank-sum test to assess differential expression of the metagene in this cluster against other clusters. To optimize hyperparameters, a subset of gene pair predictions were analyzed by a researcher who was blinded to the parameters, and who used the FCA data visualizer to rank the specificity of each gene pair in the cluster of interest, and across multiple FCA datasets. The final score for the candidate gene pair is a weighted combination of these metrics, and we output a ranked list of choices from which we select final gene pairs.

To test the efficacy of the TAB algorithm, we used it to identify a “hemocyte” cluster from the FCA 10X “Body” dataset that uniquely co-expressed the genes *Ppn* and *kuz* (**Figure 6B**). We then generated a pair of split-intein Gal4 lines designed to target the intersection of *Ppn* and *kuz* (**Figure 6B**.) We observed specific expression of *Ppn* ∩ *kuz* in larval and adult hemocytes, verified by staining with the pan-hemocyte antibody marker H2, which recognizes Hemese (**Figure 6B**) (39). The H2 antibody did not appear to stain all of the morphologically-identifiable hemocytes in the adult, consistent with previous observations (39). To test whether *Ppn* ∩ *kuz* drives expression in additional cell-types, we examined expression in sagittally-sectioned, decapitated adult flies, as well as in adult brains. In addition to specific expression in circulating hemocytes, we observed strong expression in a band of epithelial cells within the cardia, also known as the proventriculus, a structure at the foregut-midgut juncture (**Figure S5B**) (40). Interestingly, several previous studies have identified hemocyte-like at this location in the larva, which express independent hemocyte markers, and presumably play an immune function (41, 42). We confirmed that the pan-hemocyte marker *hml*-*Gal4* is expressed in a subset of the cells at this anatomical position (**Figure S5B**). Thus, our split-intein Gal4 *Ppn* ∩ *kuz* line appears to be concordant with other hemocyte-specific markers, and was not detected in other tissues.

We next generated a series of split-intein Gal4 lines to mark specific clusters based on the tissue-specific FCA “Ovary” atlas, while minimizing expression in any other cluster at the level of the whole body. We selected three clusters from the FCA ovary dataset (**Figure 6C-E**), and examined the expression of the resulting split-intein Gal4 lines *in vivo*. In all three cases, we observed the predicted expression. *CG31928* ∩ *Pez*, predicted to express in follicle cells of stage 14 oocytes, drove GFP expression specifically in these cells (**Figure 6C**). *hdc* ∩ *Nox* drove expression in the corpus luteum, a tissue composed of the follicle cells left behind after an egg is laid (43) (**Figure 6D**). *SKIP* ∩ *CG42566* drove the predicted expression in escort cells at the anterior tip of each ovariole, although we also observed expression in stalk cells between each germline cyst (**Figure 6E**).

To test whether these split-intein Gal4 lines drive expression in additional, non-targeted tissues, we examined whole adult flies, decapitated and sagittally-sectioned, as well as adult brains. Neither *CG31928* ∩ *Pez* nor *hdc* ∩ *Nox* drove detectable EGFP expression in any other adult tissue outside the desired cell type (**Figure S5C,D**). *SKIP* ∩ *CG42566* drove expression in a small number of neurons in the brain (**Figure S5E**), but was otherwise undetectable outside of the germaria, in which escort cells reside. Thus, altogether these lines were highly specific to the targeted cell type. These pilot experiments demonstrate that the TAB algorithm will be a useful tool to generate highly specific genetic tools for clusters identified by scRNAseq datasets, whether at the level of individual scRNAseq datasets, or across multiple datasets or the whole organism.

### One-step generation of double knock-in split intein Gal4 lines using dual drug-selection

One technical bottleneck in the production of split-intein Gal4 lines or split-Gal4 lines is the fact that two independent transgenic lines must be created for each desired genetic driver. We reasoned that it may be possible to generate split-intein Gal4 lines in a single step by simultaneously generating double knock-in lines. To test this approach, we adapted our knock-in vectors to contain drug selection markers that were recently characterized as transgenesis markers in *Drosophila* (44). In this approach, each knock-in is marked by a separate drug resistance gene, and double knock-in transformants are selected by rearing larvae on food containing both of the relevant drugs.

We created a modified version our Gal4^N-int^ donor vector containing a resistance gene for blasticidin (Blast^R^), and a version of our Gal4^C-int^ donor vector containing a resistance gene for G418 (G418^R^), and retained the fluorescent 3×P3-dsRed marker in both vectors. We used TAB to identify two pairs of genes marking clusters from the FCA Gut atlas: *CG13321* ∩ *CG6484* to label “enterocyte of anterior adult midgut epithelium”, and *CG14275 ∩ CG5404* to label “hindgut.” For each gene pair, we generated two separate drug-resistance knock-in vectors, with one construct resistant to blasticidin and the other resistant to G418 (**Figure 7A**).

**Figure 7.**
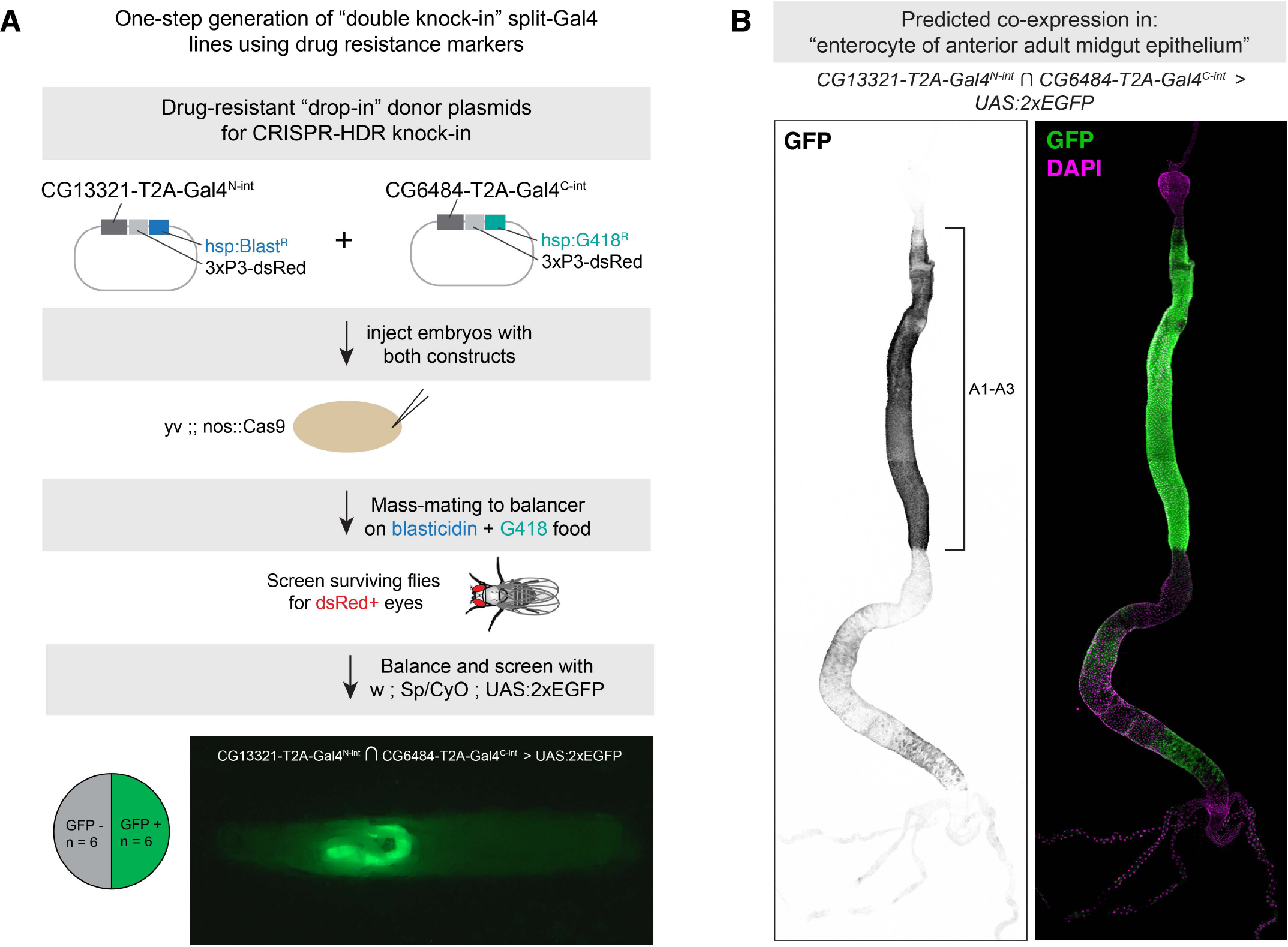
One-step generation of double knock-in split-intein Gal4 line using drug selection markers. (A) Knock-in donor vectors containing drug resistance markers (Blast^R^ or G418^R^) are co-injected into embryos expressing germline-restricted Cas9, and the offspring of these injected G0s are screened for double drug-resistance. Of the surviving F1, dsRed+ flies are screened for the ability to drive UAS:EGFP expression. Pie chart indicates the proportion of dsRed+ flies that successfully drove EGFP in the predicted cells, and image shows an L3 larva expressing EGFP in a portion of the gut, anterior to the left. (B) Expression pattern of *CG6484*-Gal4^N-int^ ∩ *CG13321-*Gal4^C-int^ in the adult gut, with anterior enterocyte regions A1-A3 (38) indicated with a bracket. Anterior is up.

We injected a 1:1 mixture of these two vectors into *nos-Cas9* embryos, and mass-mated the resulting injected G0 flies to a balancer stock, on food containing both blasticidin and G418 (**Figure 7A**). Of the flies that survived, we selected flies with *dsRed+* eyes and screened these by crossing to a UAS:2×EGFP reporter. For *CG14275 ∩ CG5404,* we only recovered a small number of flies after drug screening, zero of which were *dsRed+*. However, for *CG13321* ∩ *CG6484,* of the 12 *dsRed+* flies we screened, six (50%) drove GFP expression, indicating successful one-step creation of double knock-ins. These six lines drove strong EGFP expression throughout the anterior region of the midgut, from the posterior limit of the cardia to the anterior limit of the copper cells, as well as weaker, spotty expression in portions of the posterior midgut (**Figure 7B**.) Independently, we created separate knock-ins for *CG13321-*Gal4^N-int^ and *CG6484*-Gal4^C-int^ using our standard HDR vectors, and confirmed that these lines drove expression in an identical pattern.

We note that both of these genes are located on Chromosome 2R, indicating that it would be challenging to use standard recombination genetics to create a single chromosome containing both inserts, which further demonstrates the value of making a one-step double knock-in. However, we note an important caveat. Specifically, the double-drug selection protocol was not 100% effective, as some dsRed-negative flies were observed after the first round of mating, and 50% of the dsRed+ flies did not drive EGFP expression, indicating they were likely single knock-in flies missing one of the two selection markers. Thus, while the double knock-in strategy can serve to quickly generate split-intein Gal4 lines, it will require additional trouble-shooting to be a reliable and scalable approach.

## Discussion

In the original description of the split-Gal4 system, it was noted that the replacement of the native Gal4 AD with the VP16 activator represented a trade-off: VP16 drove much stronger expression, but rendered split-Gal4 insensitive to repression by Gal80 (8). Here, we present an alternative split-Gal4 system that obviates the need for this trade-off by generating full-length wildtype Gal4 protein from two non-functional fragments, using self splicing split inteins. The split-intein Gal4 system combines the exquisite cell type-specificity of split-Gal4 with the ability to temporally control Gal4 activity using existing Gal80^ts^ reagents. This system drives clean and specific transgene expression at similar levels to the existing split-Gal4 and Gal4 systems, and is fully repressible by Gal80^ts^. Similar to Gal4 lines, additional spatial restriction of split intein Gal4 activity should be possible using existing Gal80 lines. We believe that these advantages will make the split intein-Gal4 system a valuable addition to the toolset available to the *Drosophila* research community for targeted transgene expression in specific cell types.

Targeting of specific cell-types should be further facilitated by the widespread availability of scRNAseq datasets. As demonstrated here, such datasets can be leveraged to create intersectional split intein Gal4 tools based on knowledge of cell-type specific gene expression. This approach should allow researchers to test hypotheses generated by scRNAseq atlases, and to de-orphan clusters of unknown anatomy or function. It will also aid in the creation of highly specific drivers for nearly all cell-types and tissues in the fly, and permit functional manipulations of these cell types with temporal control. To aid in the design of cell-type specific drivers, we have developed the Two Against Background (TAB) algorithm, which we believe will reduce the potential for co-expression outside a specific cluster of interest. Future characterization of the TAB algorithm across a variety of scRNAseq datasets will help further refine the cell-type specific tools available to the *Drosophila* research community.

To facilitate the creation of split intein-Gal4 lines, we have generated a plasmid tool-kit to create split intein-Gal4 lines, either enhancer-driven or via knock-in. For knock-ins, we provide plasmids for cloning via “long” homology arms (∼1000bp), or via “drop-in cloning” using 200bp fragments, which is what we use in this manuscript. These plasmids are diagrammed in **Figure S6** and have been deposited in Addgene.

## Methods

### Experimental animals

*Drosophila melanogaster* stocks were maintained and crossed on standard laboratory cornmeal food, and experiments were conducted at 18°C, 25°C, or 29°C as indicated in the text. All adult experiments were performed in females. The new transgenic lines created in this study are described in **Table S1**, and all genotypes are provided in the Table of Genotypes below. The following previously described stocks and alleles were used:

> *tub::Gal4DBD; UAS:2×EGFP* (BL60298) - “split-Gal4 tester line”

> *tub::VP16[AD], UAS:2×EGFP* (BL60295) - “split-Gal4 tester line*”*

> *w; Sp / CyO ; UAS-2XEGFP* (BL60293)

> *w;; UAS-2XEGFP, tubGal80^ts^* (recombined from BL60293 and BL7018) esg-Gal4, UAS:GFP, tubGal80^ts^ (Perrimon lab stock)

> yki^S3A^ (BL28817)

> *w;; tub-Gal4* (Perrimon lab stock)

> *actin:GeneSwitch* (Perrimon lab stock)

> *hml-Gal4 > UAS:2XEGFP* (BL30140)

### Optimization and cloning of split-intein Gal4 components

To generate the split-intein Gal4 constructs, we proceeded in two steps. We first identified a suitable site to split the Gal4 molecule into inert fragments, and we then tested a variety of split intein pairs for their ability to reconstitute strong Gal4 activity. To generate transcriptionally inactive Gal4 fragments we sought splice sites that would divide the Gal4 DNA binding domain. We focused on serine and cysteine residues since these typically promote split intein splicing efficiency when located in the +1 position on the C-terminal side of the splice junction. We selected three cysteine residues within the Gal4 DNA binding domain (C^21^, C^28^ and C^31^) as candidate splice junctions and for each created complementary split intein Gal4 constructs using the hybrid split intein Npu DnaE_N_ and Ssp DnaE_C_ (45). To test the splicing efficacy of the three resulting pairs of split intein Gal4 constructs, we transfected S2 cells with plasmids encoding each pair and monitored the expression of a UAS-GFP reporter as described in (8). Constructs that split the Gal4 molecule at C^28^ failed to yield any Gal4 activity, whereas constructs using C^21^ and C^31^ as splice junctions were equally efficacious. The C^21^ residue was thus used as the splice junction for all further constructs. Preliminary tests *in vivo* with Npu DnaE_N_ and Ssp DnaE_C_ hybrid generated only low levels of Gal4 activity *in vivo*, so we next tested split intein Gal4 constructs using additional split inteins, including gp41-1 (19) and Cfa (46). Split intein Gal4 constructs made with gp41-1 exhibited the highest Gal4 activity and were selected for use *in vivo*.

Because GeneSwitch shares the same DNA binding domain with Gal4 (i.e. amino acids 2-93), split intein GeneSwitch constructs were generated using as the splice junction the same C^21^ cysteine residue. The split intein Gal4 and split intein GeneSwitch constructs thus share the same N-terminal Gal4[1-20] component. To create the Gal4[1-20]-gp41-1inteinN aka Gal4^N-int^ (HJP-277) and inteinC-Gal4[21-881] aka Gal4^C-int^ (HJP-287) plasmids that served as the basis for future cloning into plasmids for transgenesis, we generated synthetic oligonucleotide gBlocks (Integrated DNA Technologies, Inc., Coralville, Iowa) containing the gp41-1 and Gal4 fragments, and cloned these into expression vectors using Infusion cloning (In-Fusion HD Cloning Plus, Cat. 638911, Takara Bio USA.)

### Design and cloning of NanoTag split-Gal4 components and other constructs for experiments in S2R+ cells

All constructs for S2R+ experiments were cloned into an actin-driven expression construct derived from pAW-H2B-mCherry-12701 vector described in (22). This backbone was digested with *XbaI* and *NheI* enzymes, and various PCR-amplified inserts were then added via Gibson Assembly (New England BioLabs Cat. No. E2611.) To generate pAw-Gal4DBD-3×127D01, we first cloned Gal4DBD-1×Nb1207 by amplifying the Gal4DBD insert from pCRISPaint-T2A-Gal4-3×P3-RFP (47) and Nb127D01 from pAW-Nb127D01-GFP (22). We subsequently used Gibson Assembly to insert two additional Nb127D01 fragments. To generate pAW-NLSGal4AD-127D01, pAW-H2B-mCherry-127D01 was double-digested as described above, and the NLS-Gal4AD domain from a plasmid originally based on pActL-Gal4AD (Addgene 15303) and the 127D01 tag was inserted as part of the primer. The inserts for the original split-Gal4 components (pAW-Zip^−^-Gal4DBD and pAW-p65-Zip^+^) were amplified from pBPZpGAL4DBDUw (Addgene 26233) and pBPp65ADZpUw (Addgene 26234), respectively. pCaSpeR-tub-Gal80 was used to test for Gal80 sensitivity in S2R+ cells.

### Testing split-intein Gal4 and Nanotag split-Gal4 in S2R+ cells

*Drosophila* S2R+ cells (DGRC, 150) were cultured at 25°C, in Schneider’s media (Thermo Fisher Scientific, 21720–024) with 10% fetal bovine serum (Sigma, A3912) and 50 U/ml penicillin-streptomycin (Thermo Fisher Scientific, 15070–063). S2R+ cells were transfected using Effectene (Qiagen, 301427) following the manufacturer’s instructions. A total of 200 ng of plasmid DNA per well was transfected in 24-well plates. The cultured cells were imaged live two days after transfection on an InCell Analyzer 6000 automated confocal fluorescence microscope (GE Healthcare Lifesciences).

### Cloning and transgenesis of pTub-driven constructs

Plasmids were constructed using Gibson assembly (New England BioLabs Cat. No. E2611), as follows. A 2,595bp fragment of the *Drosophila alphaTubulin48B* promoter, followed by a 35bp minimal promoter and Kozak sequence, was PCR-amplified from pCaSpeR-tub-Gal80 (F: AAGCTTGCACAGGTCCTGTTCG R: GTTGCGGCCGCGGATCTG). This promoter was cloned upstream of either the 327bp open reading frame of Gal4^N-int^ or the 2,697bb open reading frame of Gal4^C-int^, in a backbone containing an SV40 3’UTR, an attB sequence and a mini-white marker (backbone originally from Addgene 140620) These constructs were inserted into the attP40 on the second chromosome using standard integrase-mediated transgenesis method.

### Cloning of T2A knock-in vectors via “drop-in” cloning

To generate in-frame knock-ins of split-intein Gal4 components, we followed a modification of the “drop-in” cloning technique (25). For targeting *esg, Dl,* and *Myo1A*, we designed 200bp left and right homology arms, flanking a cloning cassette containing two inverted BbsI cloning. These were synthesized and cloned into the pUC57_Kan_gw_OK backbone (25) by GeneWiz (Azenta Life Sciences). Upon *BbsI* digestion, these plasmids generate directional overhangs that allow for ligation of any insert with compatible overhangs. This backbone also includes a self-targeting sgRNA that linearizes the insert. In order to create an in-frame insert, the last nucleotide of the left homology arm was always the first nucleotide of a codon triplet in the target gene. Additionally, the last nucleotide of the left homology arm was never a thymine, which would create a TAG stop codon in our donor plasmid.

For *esg, Myo1A,* and *Dl* donor constructs, we generated drop-in-compatible split-intein Gal4 inserts via PCR. To do so, we first used Gibson assembly to subclone T2A-Gal4^N-int^ or T2A-Gal4^C-int^ into the “universal T2A HDR donor” backbone from (47), which includes a hsp70 3’-UTR and a 3XP3-dsRed transgenesis marker. We then PCR-amplified this insert (T2A-Gal4^N^ ^or^ ^C-int^-hsp70-3’UTR-3×P3-dsRed) using primers that include BsmBI sites designed to produce overhangs compatible with digested pUC57-homology arm plasmids. Following *BsmBI* digestion, these PCR fragments were ligated into BbsI-digested pUC57 donor constructs to create inserts flanked by 200bp homology arms. We also note that the Universal T2A HDR donor-Gal4^N-int^ and -Gal4^C-int^ are compatible with cloning ∼1000bp homology arms as described in (47). sgRNAs were cloned into the pCFD3 vector (48), CRISPR knock-in transgenesis was performed as previously described (49).

For knock-ins generated based on scRNAseq clusters, we streamlined the drop-in cloning process as follows. We cloned the entire T2A-Gal4^N-int^ and T2A-Gal4^C-int^ inserts, including the *hsp70-3UTR* and *3×P3-dsRed*, each flanked by inverted BsmBI sites, into the pCRBluntII-TOPO backbone, so the inserts can be released via simple digestion rather than PCR. We also switched to an updated pUC57 backbone, pUC57_Kan_gw_OK2 (50), which includes the genome-targeting sgRNA in the same backbone as the donor, so it does not need to be co-injected as an independent construct. The sgRNAs used to create these knock-in constructs are described in **Table S1**.

Drug-resistant knock-in donors were cloned using Gibson Assembly. We amplified the *hsp-BlastR* from Addgene 165911 and *hsp-G418R* from Addgene 165902 (44), with compatible overhangs to clone them downstream of the 3×P3-dsRed marker in the drop-in compatible split-intein Gal4 constructs described above.

### Generation of plasmids for enhancer-driven split-intein Gal4

To create pBP-Gal4^N-int^ and pBP-Gal4^C-int^ for LR Gateway cloning of enhancer fragments, we used Gibson Assembly to replace the Gal4DBD sequence in pBPZpGAL4DBDUw (Addgene 26233). The 2.3kb enhancer fragment VT024642 was amplified from genomic DNA using these primers: F: cctcttcaacacgcccaccaaactg, R: ctgcggctgaccacatcgagaa, with “CACC” appended to the 5’ end of the F primer to facilitate directional cloning into dTOPO-pENTR. LR Gateway cloning was conducted using standard protocols to generate VT024642-Gal4^N-int^, and standard phiC31 transgenesis was used to integrate this construct into the attP40 site.

### RU486 treatment

RU486 (Cayman Chemical Company Cat. No. 10006317) was added to standard fly food at a final concentration of 200 µm. For larval experiments, eggs were laid directly onto RU-containing food. For adult gut experiments, eggs were laid on and developed on standard food, and adults were transferred to RU-containing food for the indicated time.

### Drug selection for double knock-ins

G418 (final concentration = 250 µg/mL) and blasticidin (final concentration = 45 µg/mL) were added to 25 mL of standard food in bottles and allowed to dry, uncovered, overnight in a fume-hood. Injected flies were mass-mated to balancer lines on drug food, and flipped approximately every 3 days onto new drug-containing food. The surviving F1 offspring were screened for dsRed+ eyes, and any flies with dsRed+ eyes were then crossed to *w; Sp/CyO; 2×EGFP* to simultaneously balance and screen for double knock-ins of split intein-Gal4 components.

### Antibody staining and imaging

For sagittal sections of whole flies, decapitated adult female flies were fixed overnight in 4% paraformaldehyde, then manually sectioned using a fine razorblade (Personna by Accutec, Cat No. 74-0002). After antibody staining, bisected flies were placed in a drop of Vectashield mounting media in a 35mm, glass-bottom imaging µ-Dish (Ibidi, Cat. No. 81158.) Tissues were dissected in PBS, fixed for 20-30 minutes in 4% paraformaldehyde, and stained using standard protocols. GFP was detected using either Alexa488-coupled anti-GFP (Invitrogen A21311, used at 1:400) or chicken anti-GFP (Aves Lab GFP1020, used at 1:2000.) Hemocytes were stained using the pan-hemocyte H2 antibody (39) (Gift of Andó lab, used at 1:100.) Primary antibodies were detected with Alexa-488 or Alexa-555 coupled secondary antibodies (Molecular Probes.) Confocal imaging was performed on either a Zeiss LSM 780 or Zeiss Axio Observer Z1 with a LSM980 Scan Head, with the “Tile Scan” feature for whole guts using system defaults. Whole-larva imaging was performed on a Zeiss AxioZoom microscope. Mean pixel intensity was measured using FIJI/ImageJ, based on maximum intensity projections, with GFP+ pixels selected as regions of interest.

### Two Against Background (“TAB”) Algorithm

The selection of marker genes is a critical step in implementing the split-intein Gal4 protocol and poses unique challenges that are not addressed by existing marker-gene selection approaches. We began by articulating specific design goals for the selection algorithm: robustness to errors in scRNA-seq measurements; facilitating exploration by allowing users to choose gene pairs from a preferred set of markers; and achieving high accuracy against the whole-body background of the organism, including individual clusters in other organs. We emphasize cluster-level accuracy because high expression of the marker genes in even one off-target cluster can make *in vivo* results difficult to interpret. Existing marker-gene selection approaches are not well-suited to address these specific design objectives. Approaches that simply compare marker genes against an overall background may be unable to rule out niche cell-clusters that are a small part of the overall set, thus making the selection of precisely two markers challenging. NS-Forest v2, which was created specifically to identify a minimal set of genes to uniquely identify cell types, compares a target cluster against a set of other clusters but does not limit to marker sets with two genes. In contrast, TAB requires the identification of exactly two markers.

Our TAB algorithm addresses our design goals using an integrative approach that incorporates bulk RNA-seq data, an emphasis on selecting genes with robust expression in the cluster, and hyperparameter optimization by single-blind expert evaluations. We require the input of both the cluster of interest, as well as the anatomical tissue where this cluster resides. To ensure that the genes being selected are sufficiently highly expressed in the tissue, we examine the gene’s bulk scRNA-seq expression levels in the corresponding organ. The organ-level resolution is a valuable corrective for relatively noisy expression estimates from scRNA-seq atlases. We use the Tau statistic to quantify the specificity of any gene to the organ of interest. In addition, we minimize the dispersion (= variance/mean) that the gene’s within-cluster expression, to ensure that it is not highly variable within the cluster-of-interest.

We use an intersection of three metrics to generate a list of candidate gene pairs: the Tau and dispersion metrics, along with t-test of differentiation against all other clusters. From this candidate set, we evaluate all pairwise combinations of genes and select pairs that are effective at distinguishing the cluster of interest from others. As in NS-Forest v2, one of the metrics we consider is the number of other clusters where both the candidate genes are potential markers. We also construct a metagene as the average of the two genes, and perform the Wilcoxon rank-sum test to assess differential expression of the metagene in this cluster against other clusters. TAB outputs a ranked list of gene pairs, based on a weighted combination of these metrics.

The TAB algorithm includes a key innovation of single-blind expert-guided hyperparameter optimization to address the challenges of fine-tuning the statistical tests and metrics. With many FCA clusters being novel and poorly characterized, it is difficult to obtain ground-truth, experimentally-supported annotations of cluster markers that could be leveraged for calibrating TAB. To address this issue, we adopted an expert-guided approach: one researcher selected a set of cell clusters for evaluation, while another performed the computations for a selection of hyperparameter choices. On a subset of the clusters, the gene-pairs predictions were stripped off their associated hyperparameters and provided to the first researcher to analyze, based on inspection of the FCA data portal. The expert scores for the gene pairs were used to refine the hyperparameters, and the computational analysis was re-run on a different subset of the test cluster set. This iterative process was repeated until a satisfactory success rate was achieved. Through this single-blind expert-guided hyperparameter optimization, TAB is able to select optimal gene-pairs with high accuracy and reliability, even in the absence of ground-truth annotations.

To summarize, our TAB algorithm features several new features to the challenge of marker-gene selection. Firstly, it incorporates bulk RNA-seq data to supplement scRNA-seq expression estimates when available. Secondly, it emphasizes selecting genes with robust within-cluster expression profiles that are stable and not highly variable. Lastly, it employs an expert-guided approach for hyperparameter optimization, providing a principled and unbiased fine-tuning process.

## Acknowledgements

We thank the Microscopy Resources on the North Quad (MicRoN) core at Harvard Medical School for microscopy support. We thank Yifang Liu and Claire Hu for assistance selecting gene pairs using NS-Forest v2, and Rich Binari for assistance with fly work. The Perrimon Lab received funding from 5P41GM132087 and 5R24OD026435. The White lab received funding from the Intramural Research Program of the National Institute of Mental Health (ZIAMH002800, BHW). J.X. is supported by the start-up funding from Center for Excellence in Molecular Plant Sciences, CAS. R.S. and B.B. received funding from NIH 1R35GM141861. N.P. is a HHMI investigator. This article is subject to HHMI’s Open Access to Publications policy. HHMI lab heads have previously granted a nonexclusive CC BY 4.0 license to the public and a sublicensable license to HHMI in their research articles. Pursuant to those licenses, the author-accepted manuscript of this article can be made freely available under a CC BY 4.0 license immediately upon publication.

## Supplemental Figure Legends

**Figure S1. Pilot characterization of split-intein Gal4 and Nanotag split-Gal4 in S2R+ cells** Plasmids encoding Gal4, split-Gal4, split-intein Gal4, or Nanotag split-Gal4, each driven by a constitutively expressed Actin promoter, were transiently transfected into S2R+ cells, either with or without co-transfection of the Gal80 repressor.

**Figure S2. In the absence of *tub-*Gal80^ts^, split-intein Gal4 functions at 18°C, 25°C, and 29°C** Larvae were reared at the indicated temperature, and live-imaged at the L3 stage under identical imaging conditions. As with wildtype Gal4 (27), split-intein Gal4 activity is strongest at 29°C and decreases as the rearing temperature is lowered. The right-most panel indicates negative control, with inset showing an overexposed image to indicate the presence of larva.

**Figure S3. Enhancer-driven expression of the split-intein system** (A) Gateway LR cloning strategy for cloning split-intein Gal4 components downstream of an enhancer-of-interest. This protocol follows closely the workflow used to generate the split-Gal4 “VT” collection based on 2-3kb enhancer fragments that drive expression in the fly nervous system. (B) Proof of principle for the VT024642 enhancer fragment driving Gal4^N-int^ in the adult ISCs.

**Figure S4. Characterization of the split-intein GeneSwitch system using multiple drivers** Split-intein GeneSwitch expression in the adult ISCs using either *esg* (A) or *Dl* (B) is non-leaky, and is only observed in the presence of RU. (C) split-intein GeneSwitch expression in enterocytes throughout the adult midgut using *Myo1A*. In the absence of RU (left), expression is observed in a portion of the midgut, whereas RU drives the predicted expression throughout the gut. (D) When split-intein GeneSwitch is expressed ubiquitously using the *tub* promoter, non-RU-dependent expression is visible in portions of the hindgut. *actin*:GeneSwitch (original GeneSwitch) is expressed at lower levels than the *tub* promoter, and does not display this same leakiness. Scale bars = 50µm.

**Figure S5. Whole-body characterization of split-intein Gal4 lines generated from TAB predictions** For each split-intein Gal4 line, UAS:2XEGFP expression is shown in sagittal sections of decapitated adult female flies, as well as in the adult brain. (A) *Ppn* ∩ *kuz* expression is restricted to circulating adult hemocytes, as well as a band of epithelial cells in the cardia (also known as the proventriculus.) (B) Higher magnification of *Ppn* ∩ *kuz* in the cardia, in comparison with expression driven by the pan-hemocyte marker *hml-Gal4.* Anterior is up. (C-E) Expression of the indicated split-intein Gal4 driver lines, predicted by the TAB algorithm. In sectioned images, anterior is to the left. In brain images, dorsal is to the right.

**Figure S6. Plasmid toolkit for generating split-intein Gal4 lines via CRISPR-based knock-in or enhancer-driven** These plasmids are available via AddGene.

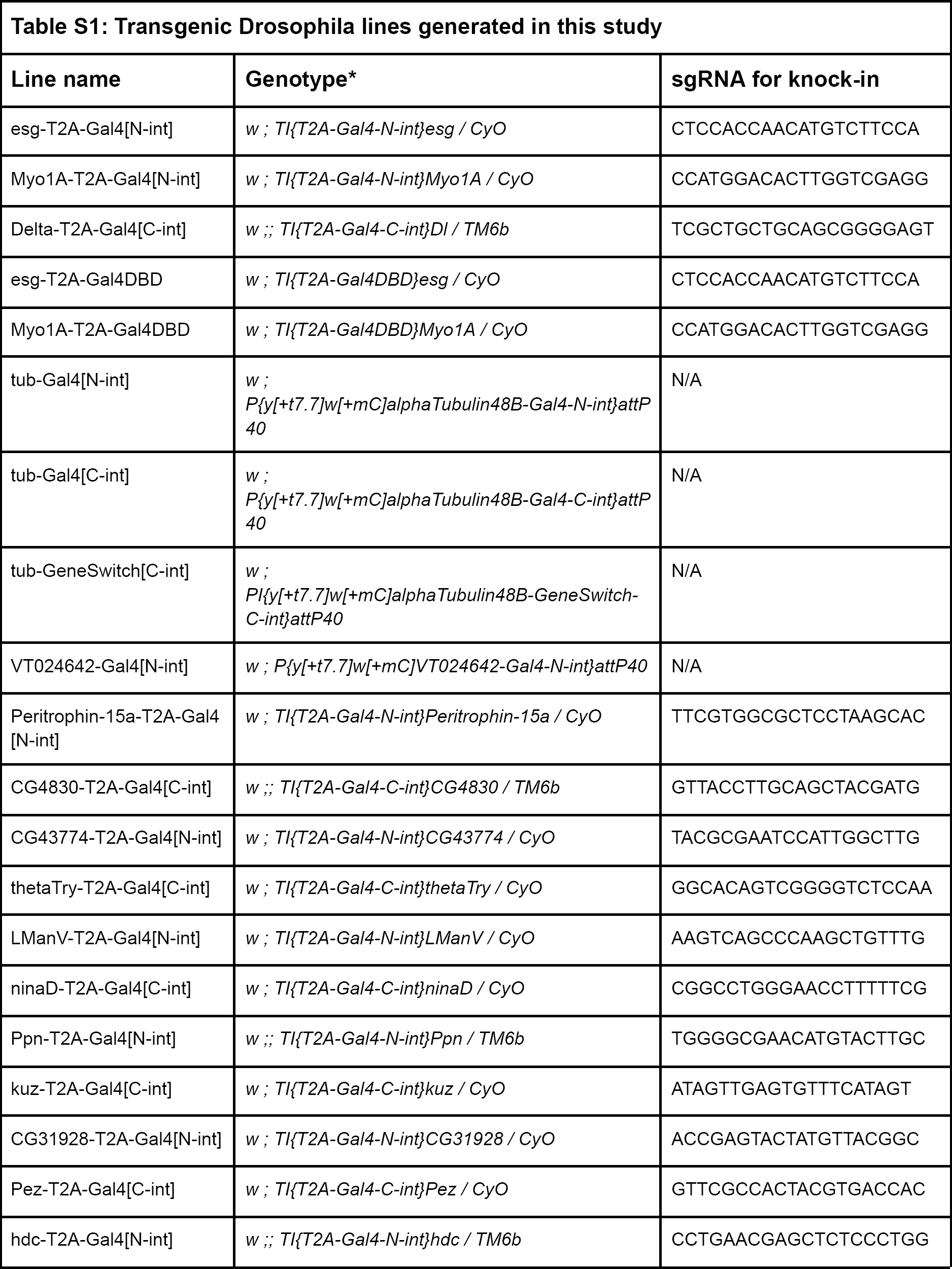
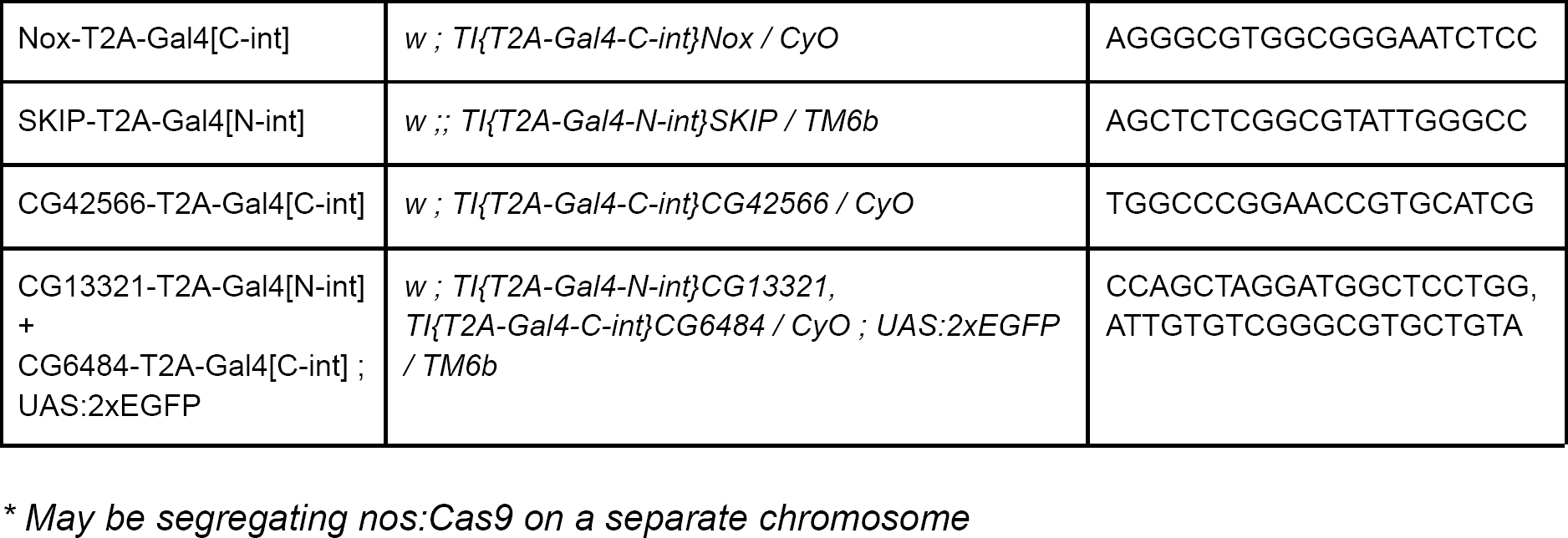

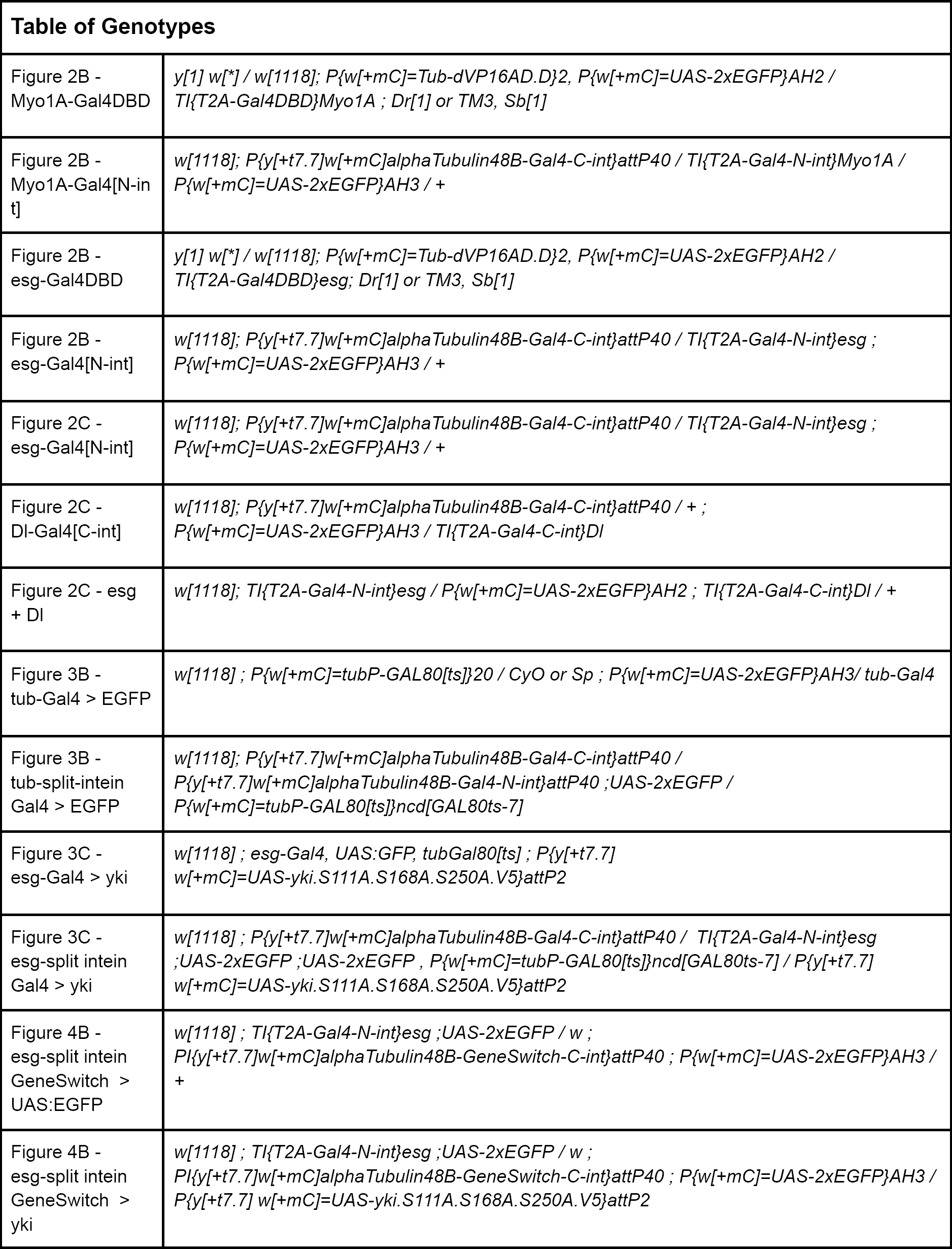
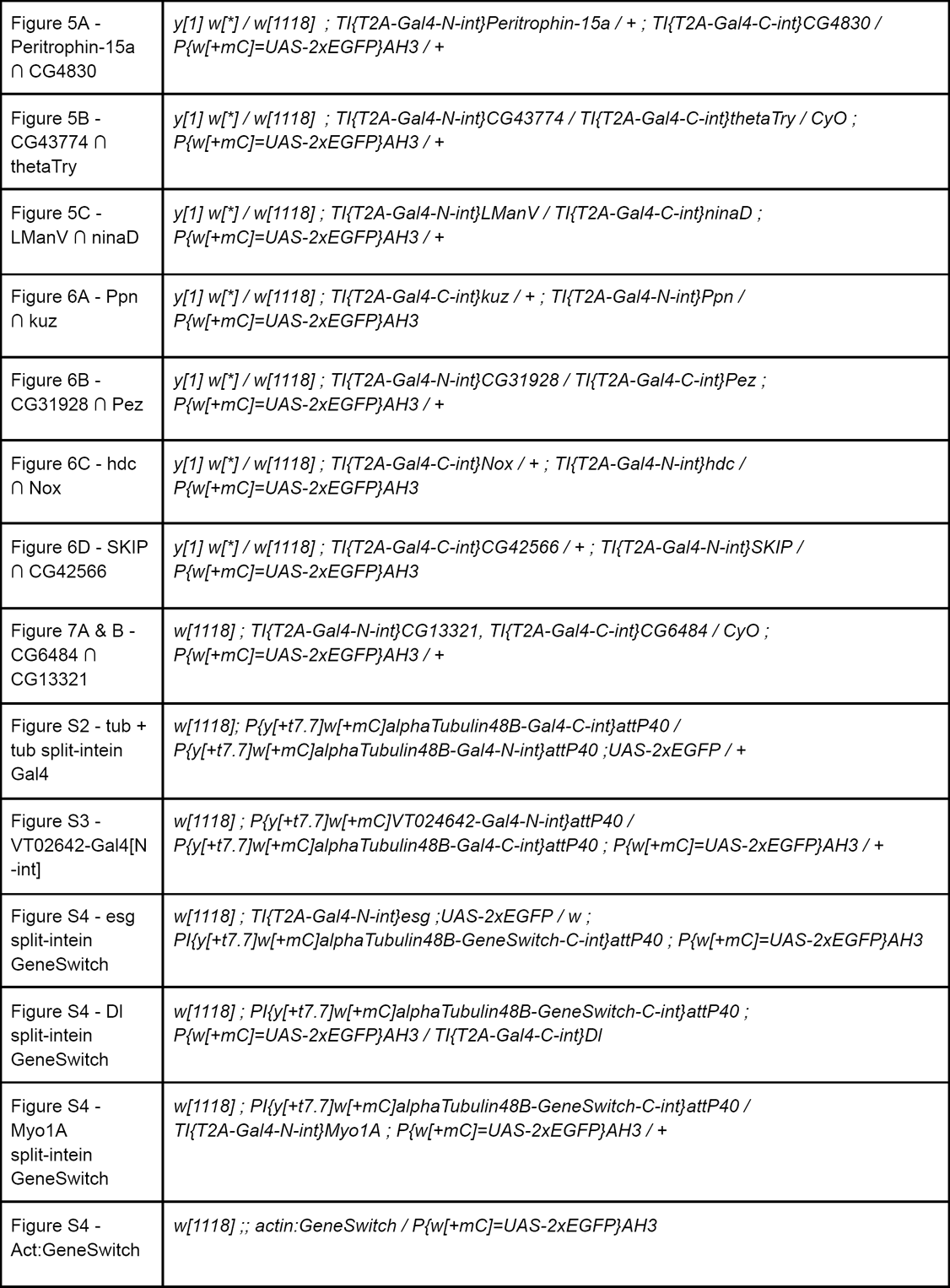
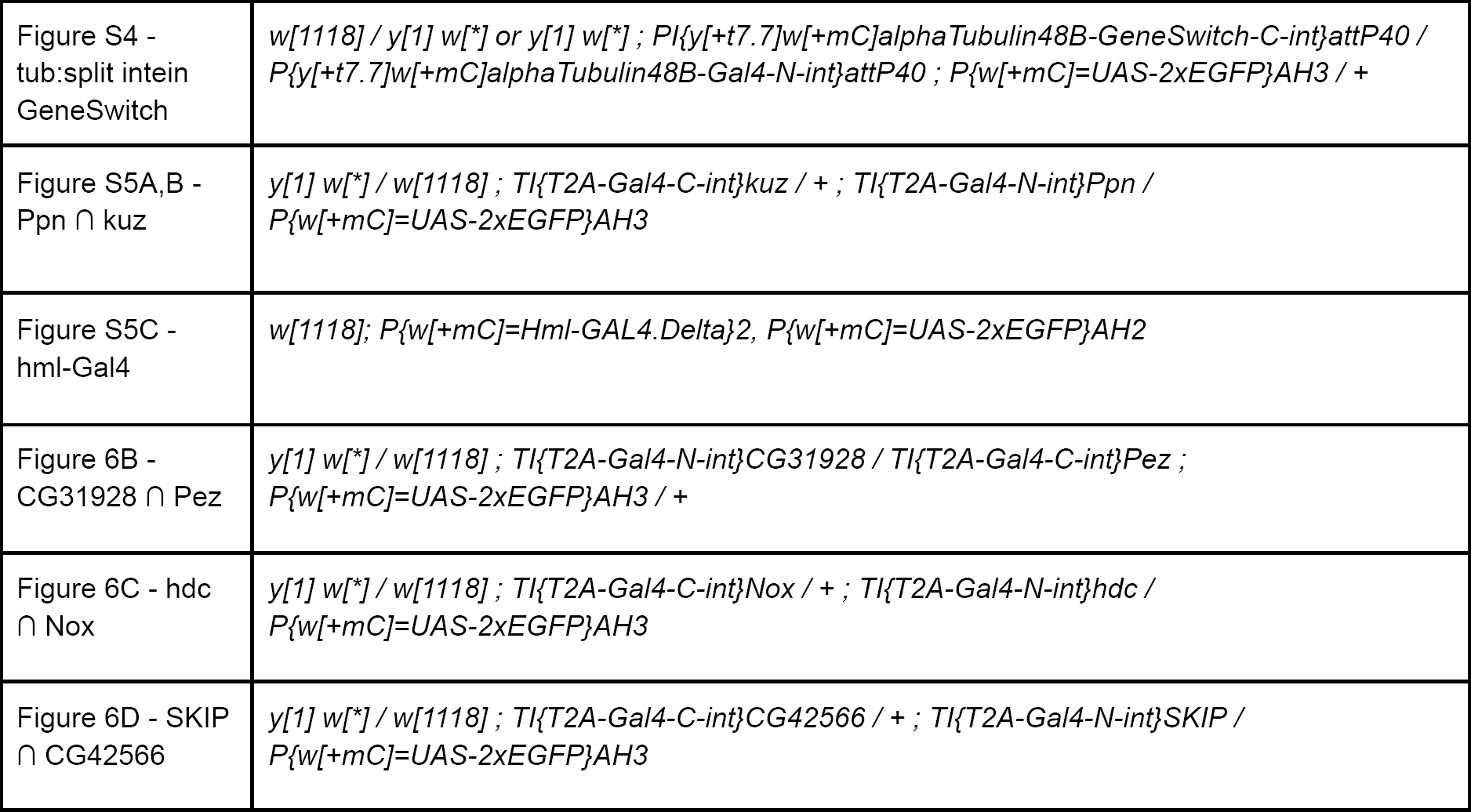

