## Supplementary material for "split-intein Gal4 provides intersectional genetic labeling that is fully repressible by Gal80": Figures S1 - S6

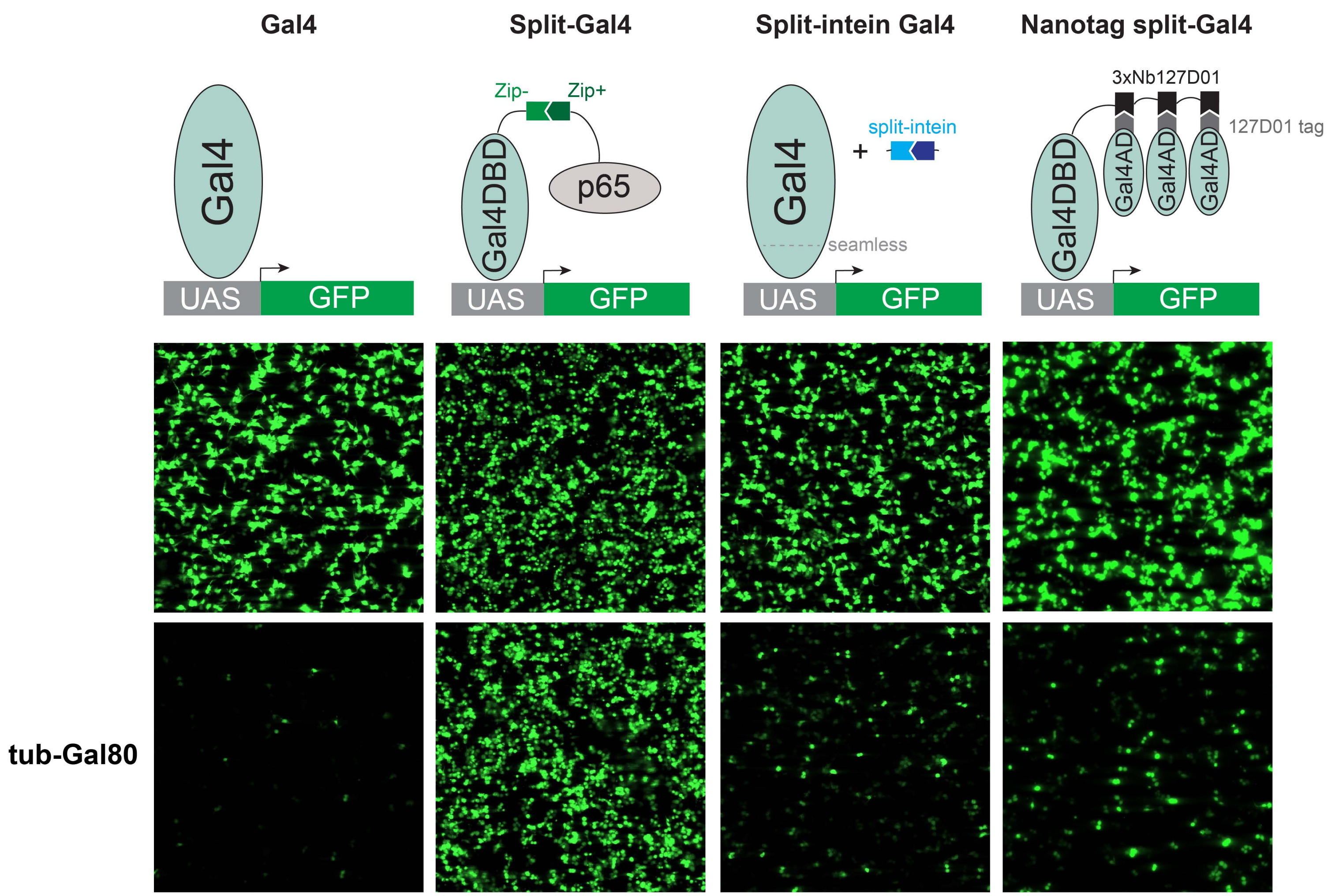
Gal4
Split-Gal4
Split-intein Gal4
Nanotag split-Gal4
Gal4
UAS
GFP
Zip-
Zip+
Gal4DBD
p65
UAS
GFP
Gal4
+
split-intein
seamless
UAS
GFP
3xNb127D01
127D01 tag
Gal4DBD
Gal4AD
Gal4AD
Gal4AD
UAS
GFP
tub-Gal80

Figure S1

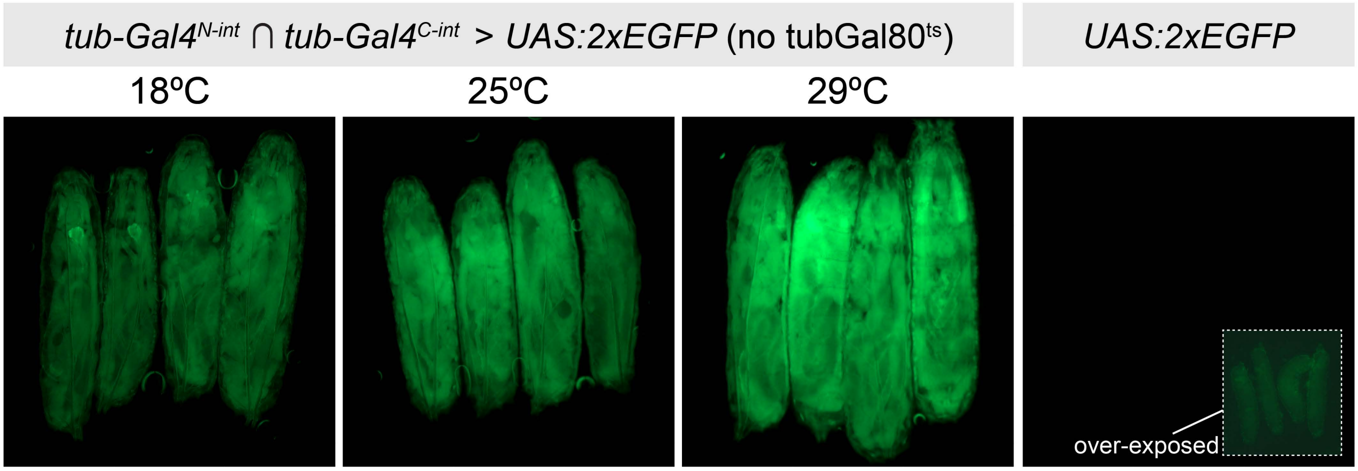

tub-Gal4^N-int ∩ tub-Gal4^C-int > UAS:2xEGFP (no tubGal80^ts)
UAS:2xEGFP
18ºC
25ºC
29ºC
over-exposed


Figure S2

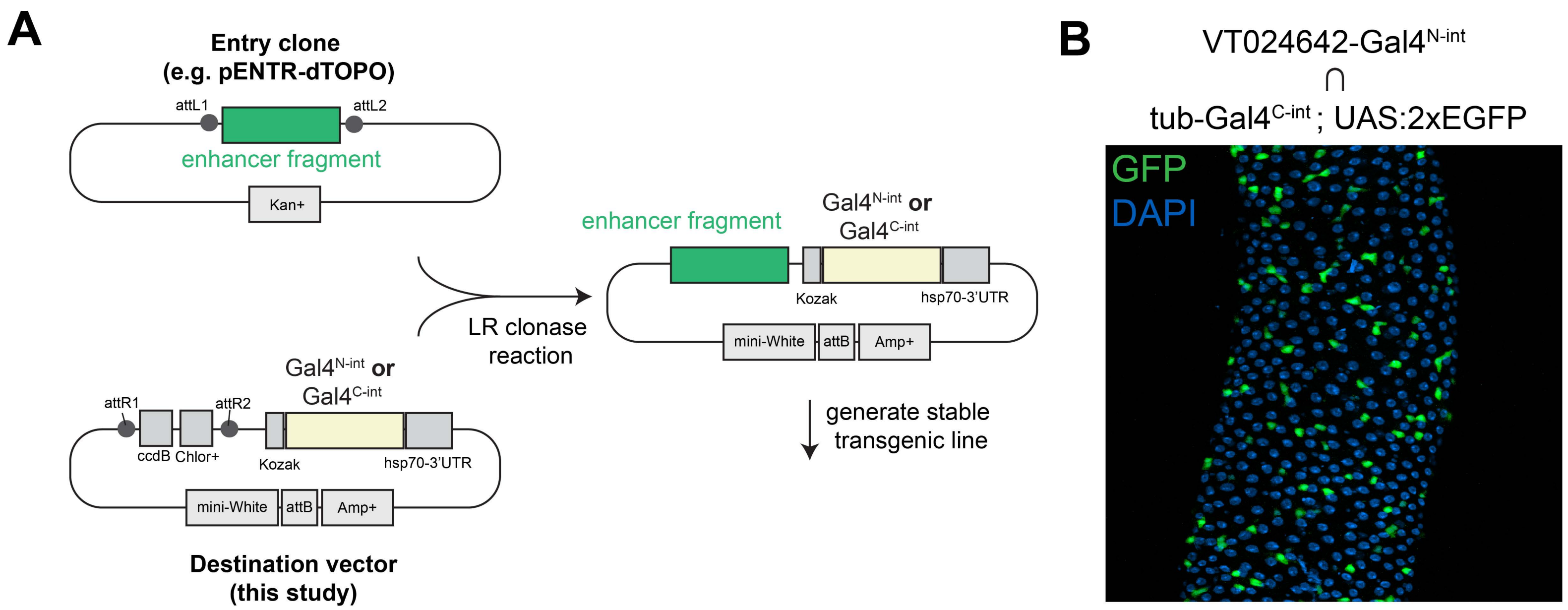
A
Entry clone
(e.g. pENTR-dTOPO)
attL1
attL2
enhancer fragment
Kan+
LR clonase
reaction
enhancer fragment
Gal4N-int or
Gal4C-int
Kozak
hsp70-3'UTR
mini-White
attB
Amp+
generate stable
transgenic line
Gal4N-int or
Gal4C-int
attR1
attR2
ccdB
Chlor+
Kozak
hsp70-3'UTR
mini-White
attB
Amp+
Destination vector
(this study)
B
VT024642-Gal4N-int
∩
tub-Gal4C-int; UAS:2xEGFP
GFP
DAPI

Figure S3

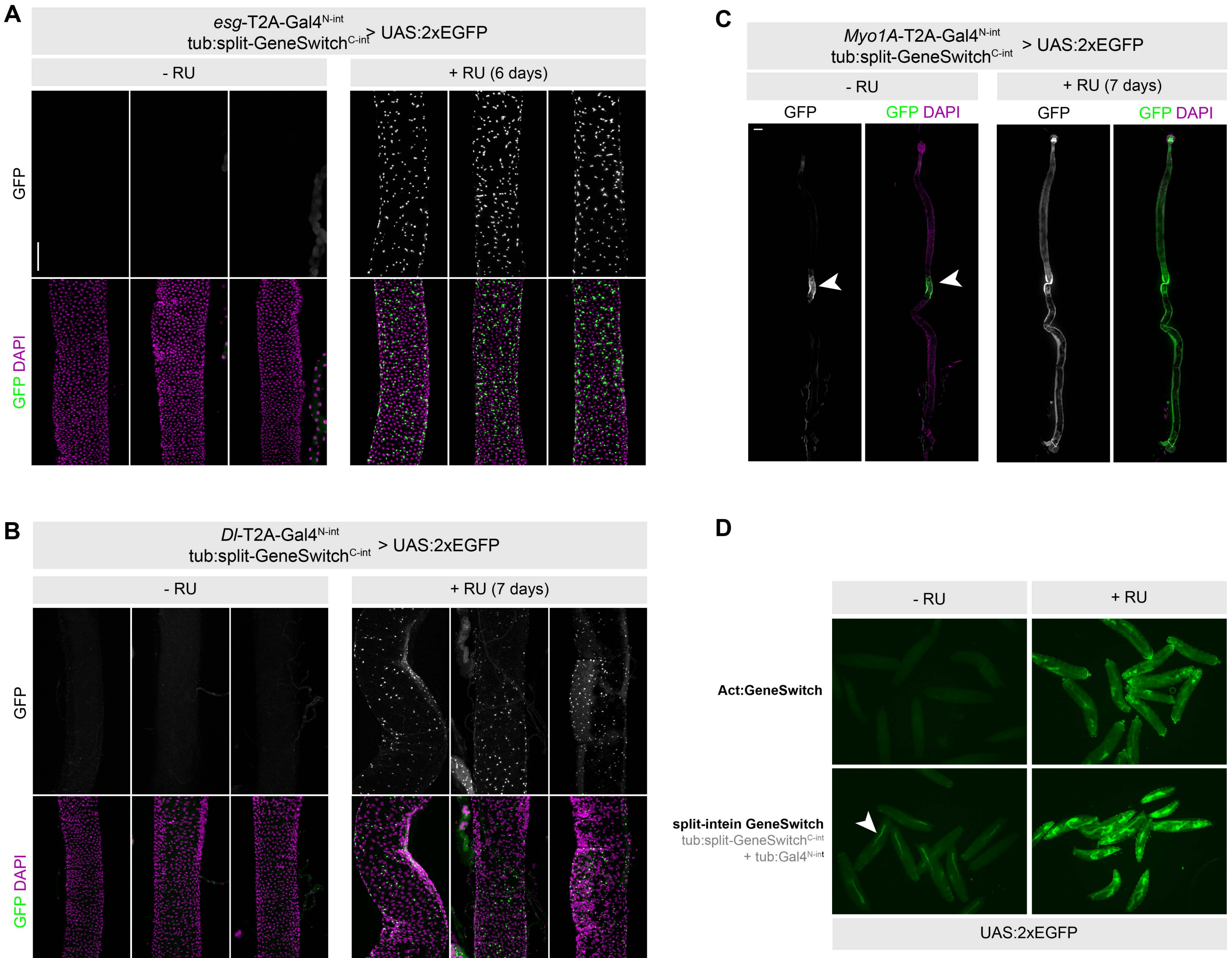
A
esg-T2A-Gal4N-int
tub:split-GeneSwitchC-int > UAS:2xEGFP
- RU
+ RU (6 days)
GFP
GFP DAPI
B
Dl-T2A-Gal4N-int
tub:split-GeneSwitchC-int > UAS:2xEGFP
- RU
+ RU (7 days)
GFP
GFP DAPI
C
Myo1A-T2A-Gal4N-int
tub:split-GeneSwitchC-int > UAS:2xEGFP
- RU
+ RU (7 days)
GFP
GFP DAPI
D
- RU
+ RU
Act:GeneSwitch
split-intein GeneSwitch
tub:split-GeneSwitchC-int
+ tub:Gal4N-int
UAS:2xEGFP

Figure S4

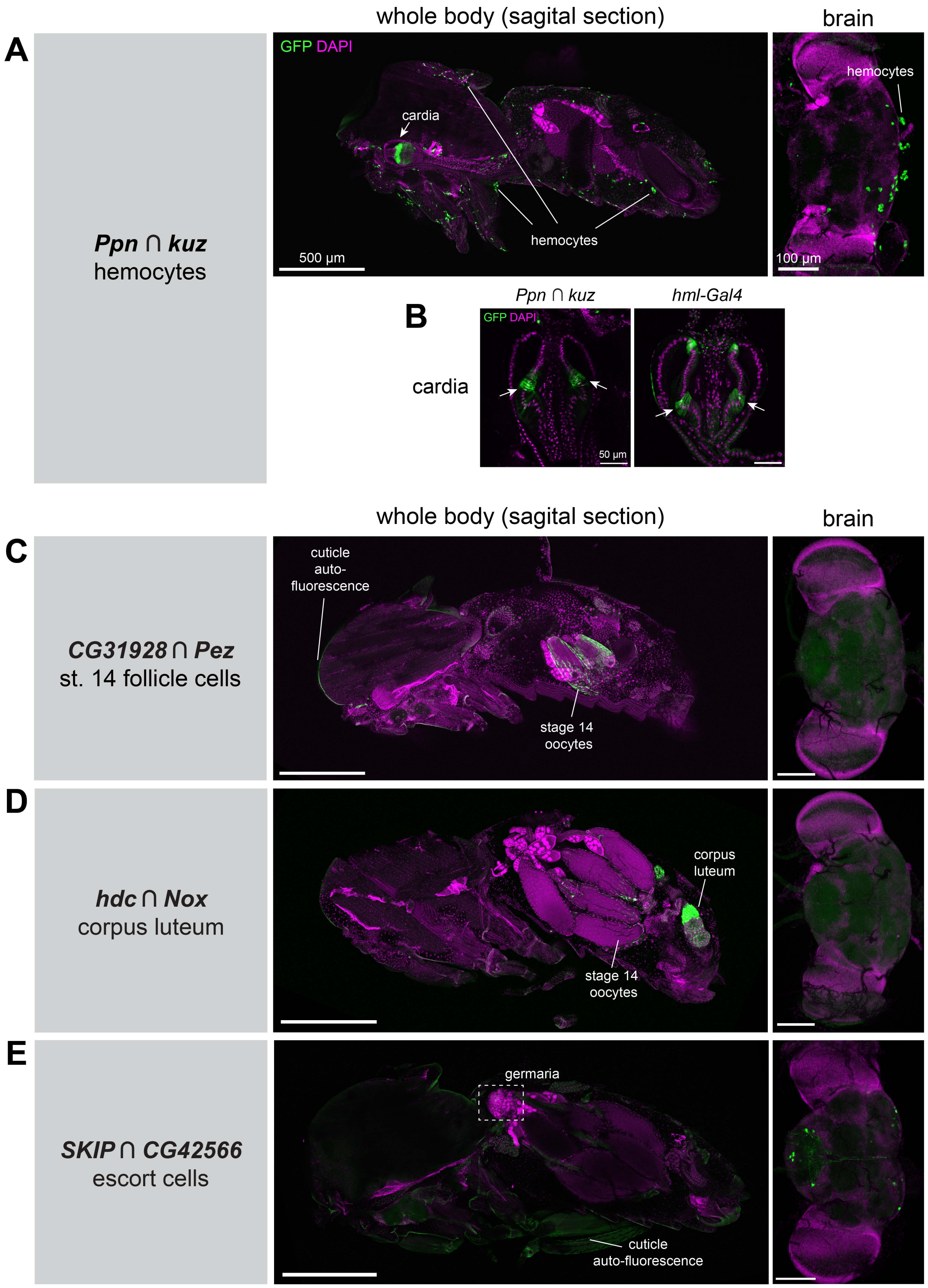

A
B
C
D
E
whole body (sagital section)
brain
GFP DAPI
cardia
hemocytes
500 µm
100 µm
***Ppn* ∩ *kuz***
hemocytes
*Ppn ∩ kuz*
*hml-Gal4*
50 µm
cuticle auto-fluorescence
stage 14 oocytes
***CG31928* ∩ *Pez***
st. 14 follicle cells
corpus luteum
***hdc* ∩ *Nox***
corpus luteum
germaria
cuticle auto-fluorescence
***SKIP* ∩ *CG42566***
escort cells


Figure S5

| Purpose (Transgenesis method) | Plasmid map | Cloning strategy |
|---|---|---|

### pHD-T2A-split-inteinGal4

Long homology arms (CRISPR knock-in)

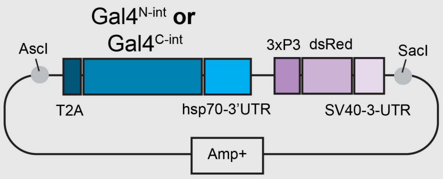


1. Digest with AscI + SacI
2. PCR amplify homology arms w/ overhangs
3. Gibson cloning

Reference: *Bosch et al. 2019*

### pDropIn-split-inteinGal4

"Drop-in" cloning (CRISPR knock-in)

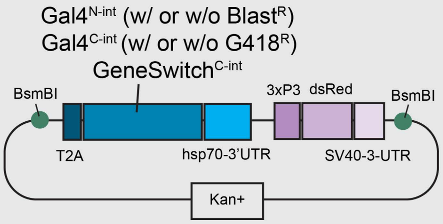


1. Release insert with BsmBI digest
2. Ligate into digested drop-in cassette

Reference: *Kanca et al. 2019*

### pBP-split-inteinGal4-destination

Enhancer driven (phiC31 integrase transgenesis)

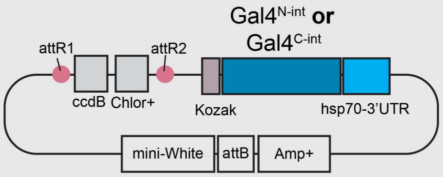


1. Clone enhancer into Gateway donor vector
2. Perform LR Clonase reaction

Reference: *Pfeiffer et al. 2008*

Figure S6
